# Pharmacological insights into safety and efficacy determinants for the development of adenosine receptor biased agonists in the treatment of heart failure

**DOI:** 10.1101/2020.07.22.215509

**Authors:** Patricia Rueda, Jon Merlin, Stefano Chimenti, Michel Feletou, Jerome Paysant, Paul J. White, Arthur Christopoulos, Patrick M. Sexton, Roger J. Summers, William N. Charman, Lauren T. May, Christopher J. Langmead

## Abstract

Adenosine A_1_ receptors (A_1_R) are a potential target for cardiac injury treatment due to their cardioprotective/antihypertrophic actions, but drug development has been hampered by on-target side effects such as bradycardia and altered renal haemodynamics. Biased agonism has emerged as an attractive mechanism for A_1_R-mediated cardioprotection that is haemodynamically safe. Here we investigate the pre-clinical pharmacology, efficacy and side-effect profile of the A_1_R agonist neladenoson, shown to be safe but ineffective in phase IIb trials for the treatment of heart failure. We compare this agent with the well-characterised, pan-adenosine receptor (AR) agonist NECA, capadenoson, and the A_1_R biased agonist VCP746, previously shown to be safe and cardioprotective in pre-clinical models of heart failure. We show that like VCP746, neladenoson is biased away from Ca^2+^ influx relative to NECA and the cAMP pathway at the A_1_R, a profile predictive of a lack of adenosine-like side effects. Additionally, neladenoson was also biased away from the MAPK pathway at the A_1_R. In contrast to VCP746, which displays more ‘adenosine-like’ signalling at the A_2B_R, neladenoson was a highly selective A_1_R agonist, with biased, weak agonism at the A_2B_R. Together these results show that unwanted haemodynamic effects of A_1_R agonists can be avoided by compounds biased away from Ca^2+^ influx relative to cAMP, relative to NECA. The failure of neladenoson to reach primary endpoints in clinical trials suggests that A_1_R-mediated cAMP inhibition may be a poor indicator of effectiveness in chronic heart failure. This study provides additional information that can aid future screening and/or design of improved AR agonists that are safe and efficacious in treating heart failure in patients.

**ONE-SENTENCE SUMMARY:** Biased agonists that preference against calcium influx relative to the cyclic AMP pathway, when compared to a conventional agonist, confer clinical safety to A_1_ adenosine receptor ligands.

## INTRODUCTION

Heart failure (HF) covers a wide range of clinical and pathophysiological conditions. It is broadly defined as a clinical syndrome whereby the heart fails to supply enough blood to fulfil the metabolic needs of the tissues (*1*). In general, the pathophysiology of HF is described by two major categories: (i) HF with reduced ejection fraction (HFrEF), where the left ventricular ejection fraction (LVEF) is <40%; and (ii) HF with preserved ejection fraction (HFpEF) where LVEF >50%. A third recently described category of HF with mid-range ejection fraction (HFmEF) is still controversial, but identifies patients with a LVEF of 40-49% in patients with different underlying characteristics and pathophysiology (*2*). There is a forecast 46% increase in HF prevalence by 2030, with over 8 million cases in the US alone (*3, 4*) illustrating the need for effective treatments.

The most recent guidelines for HFrEF treatment includes angiotensin receptor-neprilysin inhibitors (ARNI) (e.g. sacubitril/valsartan), which reduce the effect of maladaptive neurohormones and block cardiac remodelling (*5*). Although both basic research and the establishment of clear evidence-based clinical guidelines is improving management of HFrEF, these therapies have adverse haemodynamic effects (*6*). There is still a need for an approved pharmacological intervention for the treatment of HFpEF (*7*) underpinned by an increase in prevalence of this condition (*8*).

To address this, it has been suggested that therapeutic strategies should be aimed at directly attenuating adverse cardiac remodelling whilst being haemodynamically neutral (*6*). As previously suggested (*9*), fine-tuned modulation of adenosine receptors (ARs), and in particular the A_1_ subtype (A_1_R), may provide a route to fulfil these criteria, with the potential to be haemodynamically neutral, improve cardiomyocyte (CM) energetics, cardiac structure and function, and prevent further tissue injury by inducing cardioprotection and reduction of interstitial fibrosis.

Adenosine has been long-known to exert pleiotropic protective and regenerative effects and activates all four adenosine receptor (AR) subtypes (A_1_, A_2A_, A_2B_ and A_3_Rs) in different tissues (*10*). ARs are G protein-coupled receptors (GPCRs) originally classified by their pharmacological response to adenosine, with A_1_ and A_3_R inhibiting, and A_2A_ and A_2B_R activating adenylate cyclase (*11*). The adenosinergic system impacts major aspects of cardiovascular function, including beat rate, conduction, autonomic control, perfusion, growth and remodelling, and ultimately protection to injury (*12*). It is well established that all four receptor subtypes are expressed in the cardiovascular system and that their expression levels alter following injury (*13*), although full characterization of cell-specific and relative subtype abundance of AR expression is unknown.

In the setting of heart failure, ARs modulate adaptive and maladaptive responses. Cardiomyocyte hypertrophy plays an important role in this process and neurohormonal factors such as catecholamines, angiotensin II or endothelin are involved in cardiac hypertrophy and failure (*14*). Inflammation is also a hallmark of cardiac hypertrophy and involves factors such as interleukin (IL)-1β or tumor necrosis factor-alpha (TNF-α) (*15, 16*). Importantly, A_1_R agonists reduce both neurohumoral- and inflammation-driven hypertrophy (*17–19*).

Cardiac fibroblasts also contribute to the heart failure phenotype, as adverse remodelling by these cells leads to excess generation of extracellular matrix, fibrosis, and causes contractile dysfunction. The A_2B_R, which is highly expressed in fibroblasts (*20*), is the main AR subtype involved in cardiac fibroblast proliferation and collagen synthesis (*21–24*).

Despite the preclinical efficacy shown by AR agonists, further development of these agents has been compromised by the widespread expression of ARs throughout the body, and their pleiotropic effects on the cardiovascular system. These on-target side effects include modulation of blood pressure, heart rate, atrioventricular (AV) conduction, and renal function. The A_1_R (highly expressed in the atria) is responsible for changes in heart rate and conduction (*25, 26*), while the A_2A_R and A_2B_R subtypes (found in smooth muscle and endothelium) play major roles in vasoregulation (*27, 28*). Accordingly, activation of these receptors often leads to changes in blood pressure. Furthermore, AR activation plays an important role in the haemodynamic balance of the kidney, as A_1_R mediate cortical vasoconstriction and A_2A_R/A_2B_R mediate medullar vasodilation, thus reducing filtration fraction (*29*). This is an important consideration where patients with HF exhibit abnormal cardiorenal haemodynamics that ultimately exacerbate the disease (*30*).

In addition to ligands that control the strength of signalling of a GPCR (efficacy), receptors are highly dynamic proteins with different active-state conformations that can be linked to different cellular outcomes. By extension, ligands stabilising different conformations can specifically promote a subset of signalling or regulatory pathways, a phenomenon known as biased agonism (*31*). This affords the potential to target GPCRs with improved on-target specificity, as recently established in preclinical models for several GPCR agonists (*32–36*). Notably, amongst A_1_R agonists, VCP746 (and derivatives) are biased away from intracellular calcium mobilization relative to other pathways and yield more therapeutically favourable *ex vivo* pharmacology (*32, 37*).

Collectively these data suggest the possibility of identifying adenosine receptor agonists with bias profiles that yield efficacy with high therapeutic index. In 2012 Bayer described capadenoson as a non-ribose, high affinity, highly selective A_1_R agonist with good pharmacokinetics, efficacy, and a promising safety profile. It displayed reduced bradycardia in preclinical models and no effects on heart rate at rest in clinical studies (clinical study NCT00568945), while maintaining full cardioprotective potential with amelioration of markers of structural remodelling in preclinical models (*38, 39*). However, CNS safety and low solubility limited its utility and prompted the development of an improved agent in the form of neladenoson.

Neladenoson (BAY 1067197) is a pro-drug of the pharmacologically active moiety, reportedly a partial A_1_R agonist, that therefore addresses some of the limitations presented by capadenoson. Neladenoson was cardioprotective in rodents while showing fewer central side effects (*40*), and was safe and well tolerated in both phase I and phase IIa clinical studies (*41*). Despite this promising preclinical and clinical safety profile, in two phase IIb clinical trials in HFrEF and HFpEF patients (PANTHEON and PANACHE, respectively), neladenoson failed to meet its primary and secondary endpoints for efficacy (*42–44*).

Herein we sought to compare the pre-clinical pharmacology of neladenoson, capadenoson, the tool A_1_R biased agonist VCP746, and the pan-AR agonist, NECA, in molecular signalling assays across adenosine receptors, and in *in vivo* and *ex vivo* models of cardio-renal function. Neladenoson displayed high selectivity for the A_1_R, and presented a bias profile similar to VCP746 that is predictive of a lack of overt adenosine-like side effects. However, there was some divergence in other aspects of biased signalling and adenosine receptor subtype activity that might help in the design of future agents that are not only safe, but which have efficacy in treating heart failure in patients.

## RESULTS

### Neladenoson and capadenoson are differentially biased at the adenosine A_1_ receptor compared with VCP746

In stably-transfected CHO-A_1_R cells, VCP746, neladenoson, and capadenoson were all partial agonists for calcium mobilisation relative to the prototypical full agonist NECA (Fig 1A), yet displayed full agonism for inhibition of cAMP accumulation (Fig 1B). The calcium response was sensitive to pre-treatment with pertussis toxin (*data not shown*), suggesting that, like cAMP inhibition and ERK1/2 phosphorylation (*45*), it is downstream of canonical Gα_i/o_-coupling. Analysis of bias, using NECA as a reference agonist and cAMP inhibition as the reference pathway, demonstrated that the synthetic ligands were biased *away* from calcium mobilisation relative to cAMP (Fig 1C), a property that is reported to be predictive of a lower propensity for bradycardia.

**Figure 1.**
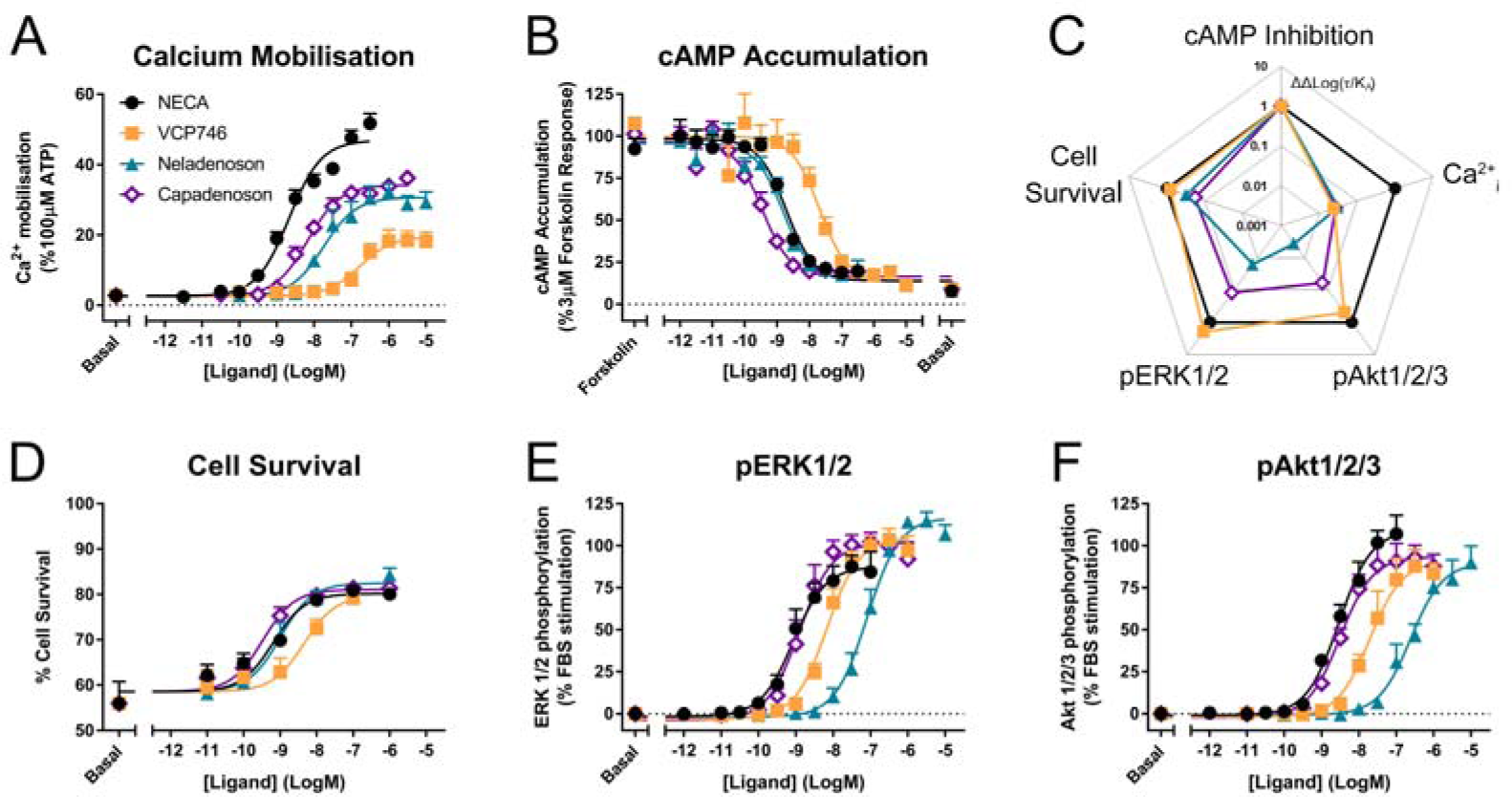
Multipathway profiling at the A_1_R. Adenosine receptor agonists were analysed in CHO-hA_1_R cells in assays of calcium mobilisation (A; n=4), inhibition of cAMP accumulation (B; n=6), cell survival (D; n=3) or phosphorylation of ERK1/2 (E; n=5) and Akt1/2/3 (F; n=3). These data were used to calculate bias, where Log(τ/K_A_) are normalised to NECA (ΔLog(τ/K_A_)) and cAMP accumulation (ΔΔLog(τ/K_A_); C). Statistical significance of bias is reported in Supp Figure 1. Concentration response data are expressed as mean ± SEM.

However, neladenoson and capadenoson differed from the VCP746 profile for cell survival (1D), and phosphorylation of ERK1/2 (Fig 1E) and Akt1/2/3 (Fig 1F; Supp Table 1), being biased away from these endpoints compared to NECA and VCP746 (Fig 1C; Supplementary Fig 1). Interestingly, despite the structural similarity of the two investigational agents, neladenoson consistently displayed lower potency than capadenoson across all assays, particularly for protein phosphorylation (Figs 1E, 1F). Neladenoson, like VCP746, is biased away from Ca^2+^ influx relative to the cAMP pathway, a profile linked to reduced adenosine-like side effects. However, unlike VCP746, neladenoson shows additional bias away from the MAPK pathway.

### Functional adenosine receptor subtype selectivity and differential biased agonism at the adenosine A_2B_ receptor

As all adenosine receptor subtypes potentially contribute to the *in vivo* activity of AR agonists, the ligands were evaluated in CHO cells stably expressing the A_2A_R, A_2B_R or A_3_R subtypes. Compared to the reference agonist NECA, VCP746 had high potency at the A_2B_R and weak activity at the A_2A_R and little or no activity at the A_3_R (Fig 2; Supplementary Fig 2; Supplementary Tables 2-4). Capadenoson stimulated other AR subtypes with an order of selectivity of A_1_ > A_2B_ > A_2A_ >> A_3_. In contrast, neladenoson had no measurable activity at the A_2A_R or A_3_R, but activated the A_2B_R as a partial, biased agonist except for cAMP inhibition where it was a low potency full agonist (Fig 2; Supplementary Tables 2-4).

**Figure 2.**
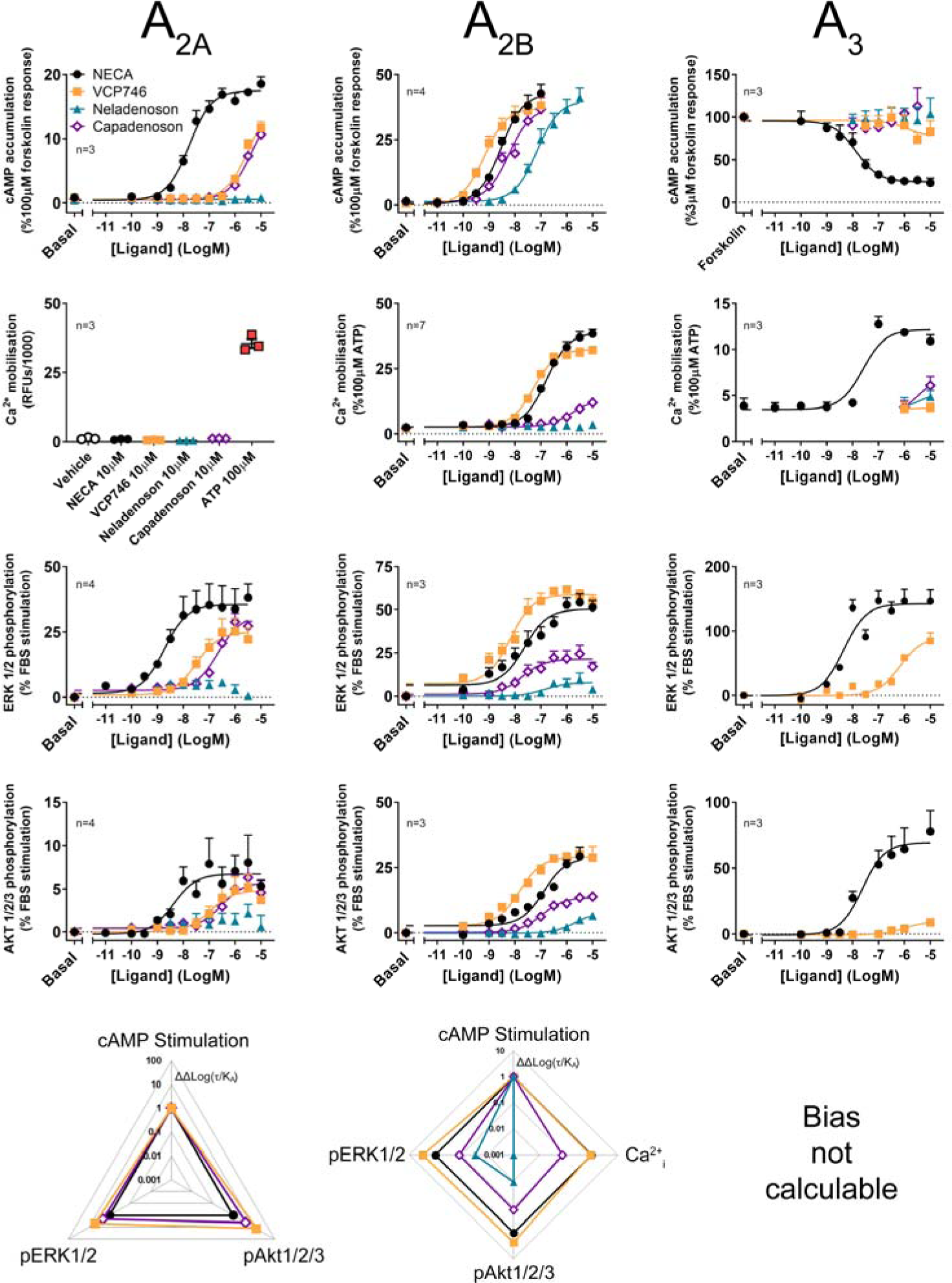
Multipathway profiling at the A_2A_R, A_2B_R and A_3_R. Adenosine receptor agonism was analysed in CHO-hA_2A_R (left panels), CHO-hA_2B_R (centre panels) and CHO-hA_3_R cells (right panels). Cells were analysed in cAMP (top row), calcium (2^nd^ row), and phosphorylation of ERK1/2 (3^rd^ row) and Akt1/2/3 (4^th^ row), with calculated bias shown in the lower row. As bias data are normalised to NECA and the cAMP pathway, where there is no response to NECA at a pathway, ligand-induced cAMP in a cell line, bias cannot be calculated. Statistical analyses of bias are shown in Supp Figure 1. Concentration response data are expressed as mean ± SEM.

The signalling profile of neladenoson, capadenoson, and VCP746 at the A_2B_R revealed marked differences (Fig 2). In contrast to the A_1_R, VCP746 was a non-biased full agonist at the A_2B_R, with NECA-like activity for inhibition of cAMP accumulation, calcium mobilisation, ERK1/2, and Akt1/2/3 phosphorylation. On the other hand, despite being a full agonist for cAMP accumulation at the A_2B_R, capadenoson was a weak, partial agonist, biased away from calcium mobilisation, ERK1/2, and Akt1/2/3 phosphorylation relative to NECA. As with the A_1_R, neladenoson was less potent than capadenoson at the A_2B_R subtype, showing bias away from ERK1/2 and Akt1/2/3 phosphorylation relative to NECA, and had no detectable activity on calcium mobilisation (Fig 2; Supplementary Table 3; Supplementary Fig 1).

Agonist activity at the A_2A_R was also different for neladenoson. In contrast to capadenoson and VCP746, which were weak, partial, non-biased agonists, neladenoson did not activate the A_2A_R subtype (Fig 2; Supplementary Table 2). Of the test agents, only VCP746 showed agonist activity at the A_3_R (in pERK1/2; Fig 2), and was a low affinity antagonist for A_3_R-stimulated calcium mobilisation and pERK1/2 (Supplementary Fig 2). Taken together, the pharmacological characterisation of these compounds at adenosine receptor subtypes shows that neladenoson is a selective A_1_R biased agonist, with biased, weak agonism at the A_2B_R subtype, while VCP746 is a biased A_1_R agonist and potent unbiased agonist at A_2B_R.

In order to broadly assess the physiological and pathophysiological effects of VCP746 and neladenoson in heart failure, these compounds were examined in a range of *ex vivo* and *in vivo* models to study beat/heart rate, hypertrophy/remodelling, aortic vasorelaxation, and renal vasoconstriction and vasorelaxation (Fig 3).

**Figure 3.**
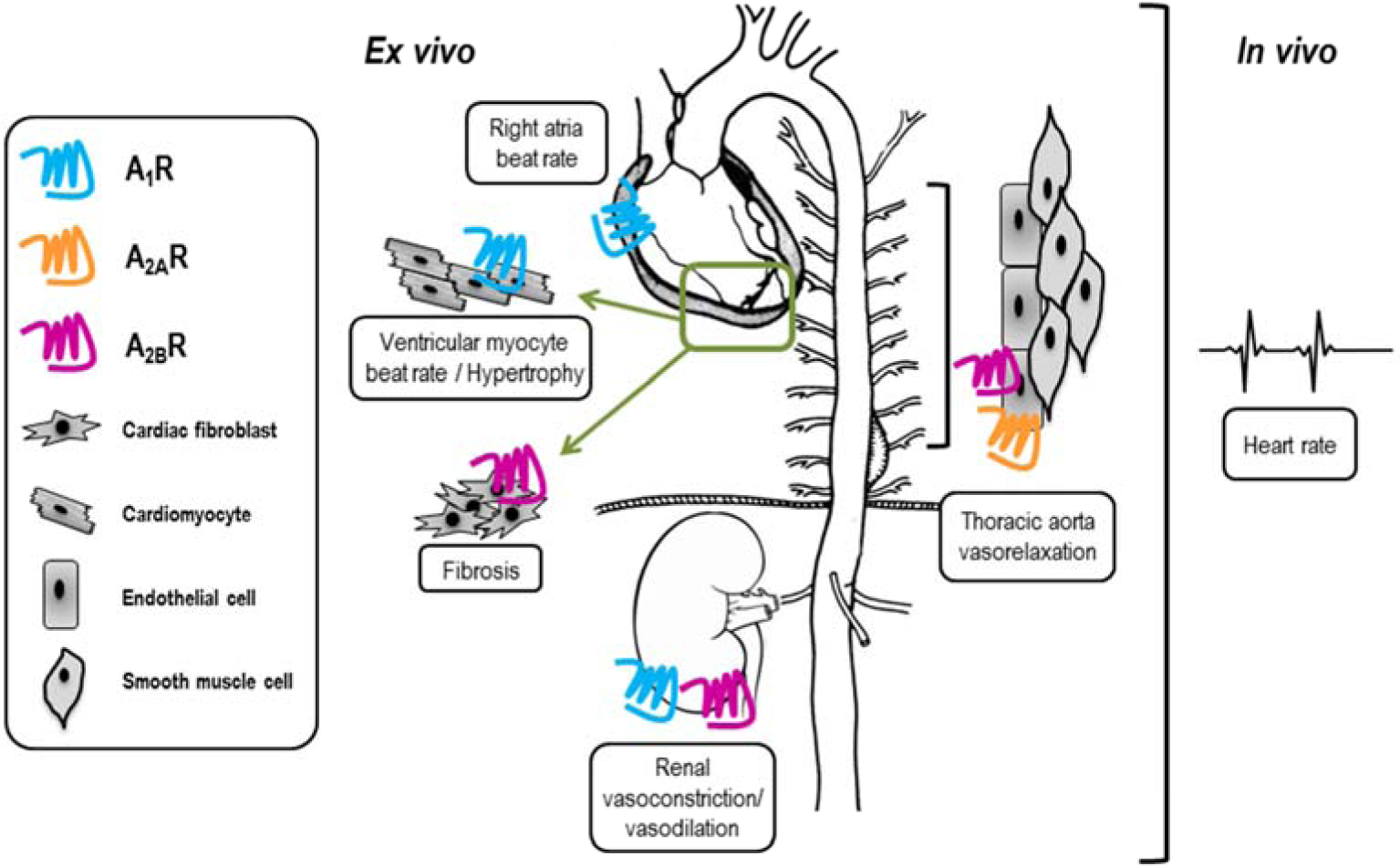
Tissue-specific expression of adenosine receptors may influence side-effect profile of compounds used *in vivo*. VCP746 and neladenoson were studied further in the indicated *ex vivo* and *in vivo* assays (boxed) to determine expected side-effect profiles. Known sites of adenosine receptor expression are indicated by blue (A_1_R), orange (A_2A_R) and pink (A_2B_R) receptors.

### VCP746 and neladenoson are anti-hypertrophic in cardiomyocytes

Putative anti-remodelling effects were assessed by [^3^H]-leucine incorporation in cardiac myocytes as a marker of hypertrophy and remodelling associated with chronic heart failure. The anti-hypertrophic effect of VCP746 is believed to be mediated by A_1_R (*17*). In the current study TNFα, IL1β, or Ang II induced neonatal ventricular cardiomyocyte (NVCM) hypertrophy that was prevented by pre-treatment with VCP746 or neladenoson in a concentration-dependent manner (Fig 4). These effects occurred over the same range of concentrations for both drugs, despite the 10-fold higher potency of neladenoson for A_1_R mediated cAMP inhibition (Fig 1). None of the compounds showed a significant effect on NVCM viability as measured by both PI staining or LDH release assays (Supplementary Fig 3), indicating that the inhibitory effects are not due to a reduction on cell viability.

**Figure 4.**
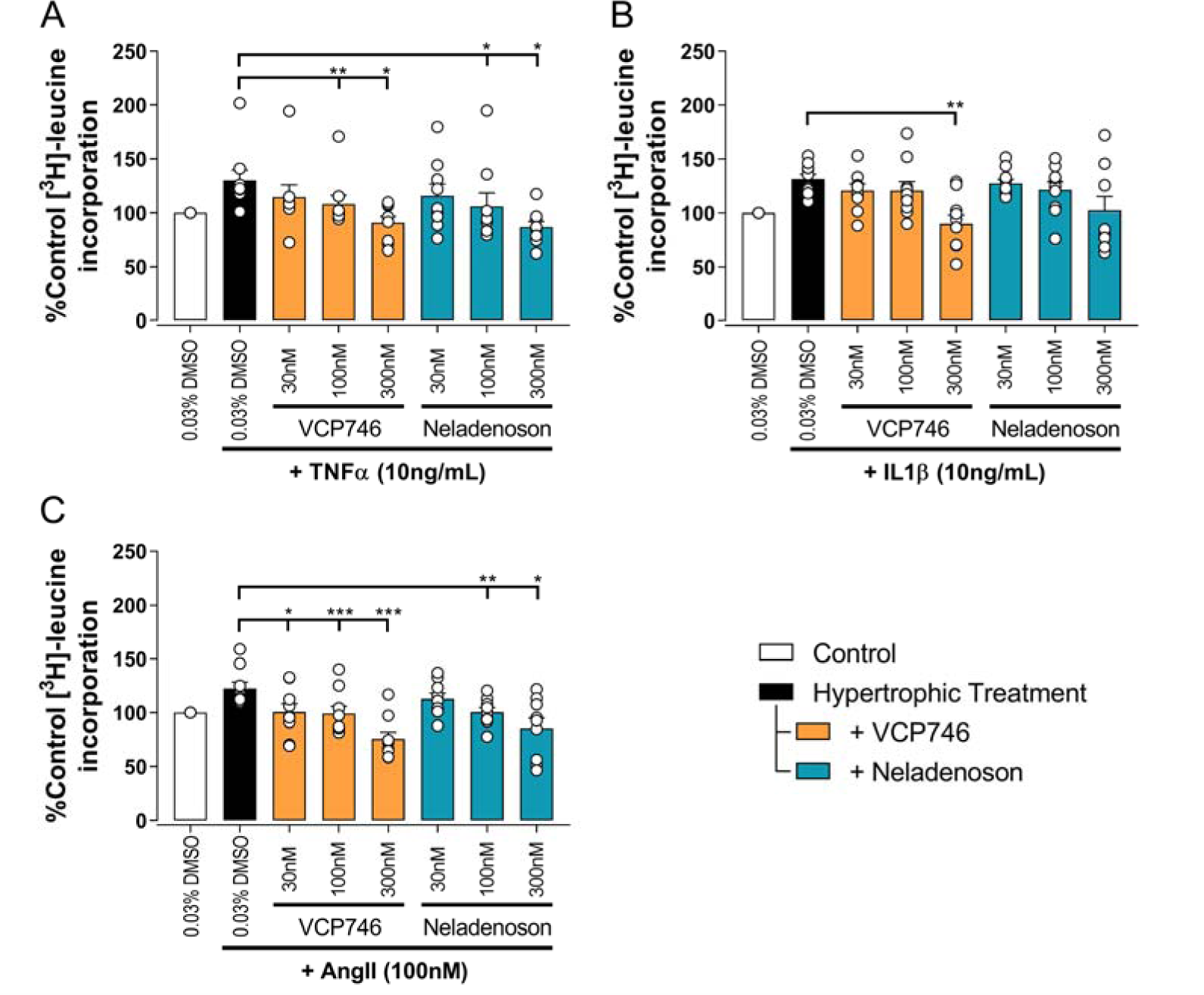
VCP746 and neladenoson reduce cardiomyocyte hypertrophy. Rat primary neonatal ventricular cardiomyocytes were exposed to hypertrophic stimuli TNFα (10ng/mL; A; n=9), IL1β (10ng/mL; B; n=10) or AngII (100nM; C; n=9) for 72h after 2h pretreatment with the indicated adenosine receptor agonists. Hypertrophy was assessed by incorporation of ^3^H-leucine as an indicator of protein synthesis. Statistical significance was assessed by repeated measures one-way ANOVA with Dunett’s post-test, compared to hypertrophic treatment (black bar) alone (*p<0.05, **p<0.01, ***p<0.001).

### Both VCP746 and neladenoson have anti-fibrotic effects in cardiac fibroblasts

In cardiac fibroblasts, Ang II and tumour growth factor-beta (TGFβ) significantly increased [^3^H]-proline incorporation, a marker of fibrosis (Fig 5). Pre-treatment with either VCP746 or neladenoson (300 nM) reduced the effect of Ang II by 40% (Fig 5A), putatively by activation of adenosine A_2B_ receptors (*46*), although neither agent affected [^3^H]-proline incorporation driven by TGFβ (Fig 5B). These effects are consistent with the signalling profile of the agonists at the A_2B_R since both are able to activate this receptor.

**Figure 5.**
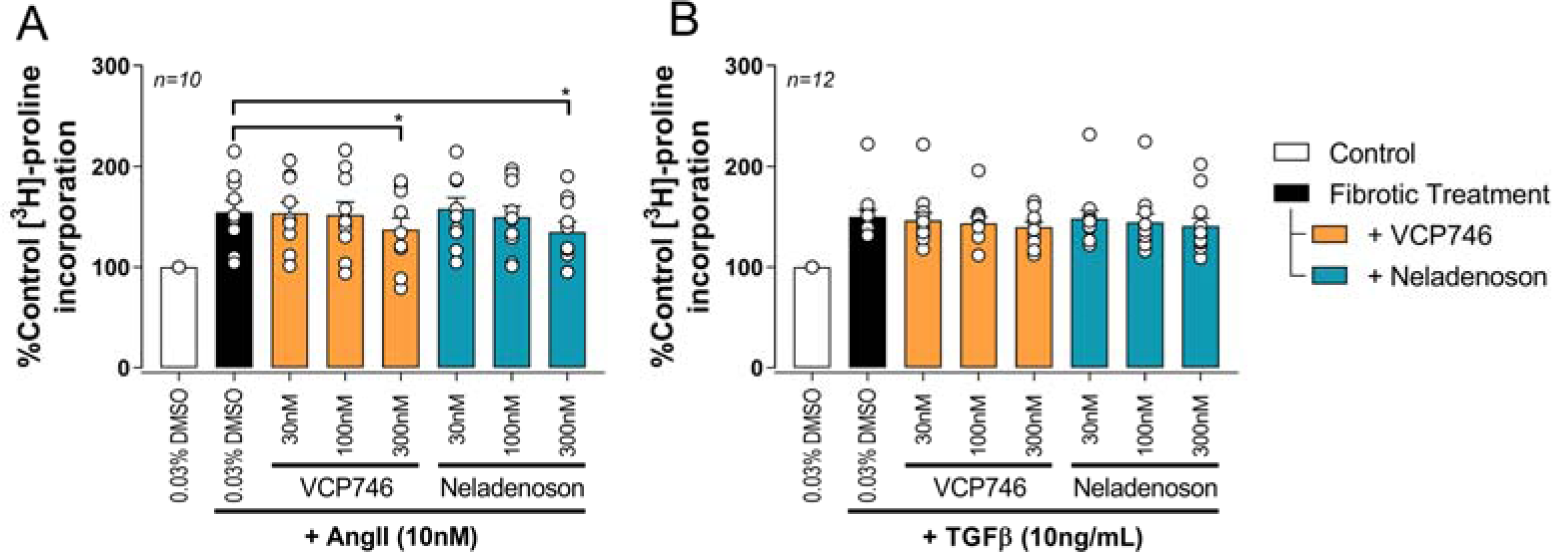
Adenosinergic agonists show some anti-fibrotic activity in rat primary neonatal cardiac fibroblasts. Fibrosis was measured by the incorporation of ^3^H-proline as an indicator of collagen formation. After 72h 10nM AngII (A) or 10ng/mL TGFβ (B) treatment, VCP746 and neladenoson pre-treatment (2h) inhibited the AngII-fibrotic effect only, at 300nM (repeated measures one-way ANOVA, Dunnett’s post-test, in comparison to fibrotic treatment alone; *(p<0.05)).

### VCP746 and neladenoson display limited and differential effects on beat / heart rate *in vitro* and *in vivo*

One of the major barriers to the use of adenosine A_1_R agonists in the clinic is A_1_R-mediated negative chronotropy i.e. bradycardia. To evaluate the relative propensity to modulate beat rate, we compared VCP746 and neladenoson head-to-head in primary NVCMs, rat atria *ex vivo*, and in conscious rats by telemetry.

In primary rat NVCMs, neladenoson (30 and 300 nM) produced a small, but significant reduction in spontaneous beat rate (Fig 6A), whereas VCP746 (300 nM) was without significant effect. The reduction of beat rate after treatment with neladenoson was reversed by pre-treatment with an A_1_R subtype-selective antagonist, SLV320, indicating that this effect is A_1_R-mediated (Fig 6A).

**Figure 6.**
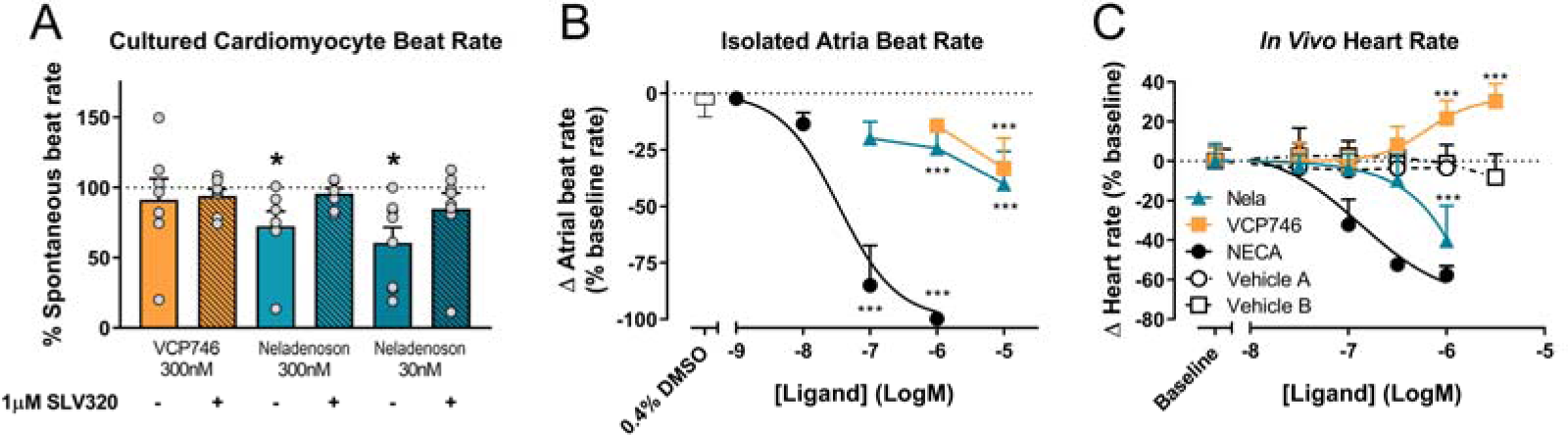
*Ex vivo* and *in vivo* A_1_R-mediated chronotropic effects. In isolated neonatal ventricular cardiomyocytes, VCP746 (n=7) had no effect on the spontaneous contraction rate of cells (A), while neladenoson (30nM: n=7; 300nM: n=8) showed a significant depressive effect, which is completely abrogated by pre-treatment with the A_1_R antagonist SLV320 (statistics were performed on raw data; paired student’s t-test, comparison of VCP746/neladenoson-treated cells to the same cells prior to treatment; *(p<0.05)). Data are mean ± SEM. In isolated atria (B), NECA produced a concentration-dependent effect (n=8), while only minor effects of VCP746/neladenoson (n=4-9) were observed at high concentrations (1 or 10μM; one-way ANOVA compared to vehicle alone; ***(p<0.01)). (C) Adenosinergic compounds were also tested *in vivo*. NECA (n=6) decreased heart rate relative to vehicle A (n=2). Neladenoson (n=4) significantly reduced heart rate at 1µM relative to vehicle B (n=7), while VCP746 (n=5) increased heart rate at high concentrations *in vivo* (two-way ANOVA, Sidak’s post-test, compared to vehicle B). Data for (B) and (C) are expressed as mean ± SD.

In rat isolated right atria, neladenoson and VCP746 had a minimal effect on the beat rate in contrast to NECA, which decreased beat rate in a concentration-dependent manner (pIC_50_ = 7.5 ± 0.3, n=8; Fig 6B). By using AR subtype-specific agonists and the A_1_R specific antagonist SLV320 we confirmed that the bradycardic response was also almost entirely A_1_R-mediated. The A_1_R selective agonist 2-Me-CCPA maximally inhibited beat rate in a manner similar to the non-selective agonist, NECA, whereas CGS21680 (A_2A_R selective) or BAY60-6583 (A_2B_R selective) had no significant effect in the same concentration range (Supplementary Fig 4A). Additionally, pre-treatment with the selective A_1_R antagonist SLV320, abrogated the chronotropic effect of NECA, further confirming the involvement of A_1_R (Supplementary Fig 4B).

Likewise, in conscious rats, increasing plasma concentrations of NECA reduced telemetered heart rate in a concentration-dependent manner (pIC_50_ = 6.9 ± 0.2, n=6; Fig 6C). Neladenoson (1 μM) produced a decrease in heart rate, whereas VCP746 (1 - 3 μM) produced a modest, concentration-dependent increase in heart rate (Fig 6C).

Collectively the data show that AR-mediated chronotropic effects are A_1_R-dependent and suggest that, unlike prototypical AR agonists, biased agonists such as VCP746 and neladenoson have limited effects on cardiomyocyte or isolated atria beat rate *ex vivo,* and heart rate in rodents *in vivo*, further strengthening the link between the bias profile of both of these compounds and the lack of chronotropic effects, thus providing a potential route for avoidance of on-target mediated side effects.

### VCP746 and neladenoson induce renal vasodilation by A_2B_R agonist activity

Patients with heart failure exhibit abnormal cardio-renal haemodynamics that ultimately exacerbates the disease. Adenosine receptors play an important role in the haemodynamic balance in the kidney; adenosine induces A_1_R-mediated cortical vasoconstriction and A_2A_R/A_2B_R-mediated medullar vasodilation, reducing filtration fraction in order to recover from negative energy balance in the kidney. Accordingly, effects in the kidney may potentially have implications for any adenosine receptor agonist synthesised for clinical use.

Using the α_1_-adrenoceptor agonist methoxamine to elevate vascular tone, we investigated renal vasodilation. The adenosine A_2B_R agonist BAY 60-6583, VCP746, and neladenoson all induced vasodilation, with a potency (pEC_50_) of order VCP746 (10.1 ± 0.3, n=3) > BAY 60-6583 (9.4 ± 0.4, n=3) > neladenoson (8.4 ± 0.2, n=3; Fig 7A). Since the BAY 60-6583 response is sensitive to MRS1754, but not SCH442426 (Supplementary Fig 5A), it suggests that the response is A_2B_R-, rather than A_2A_R-mediated. This would be consistent with VCP746 showing greater potency and efficacy at the A_2B_R compared with neladenoson (Fig 7A; Fig 2).

**Figure 7.**
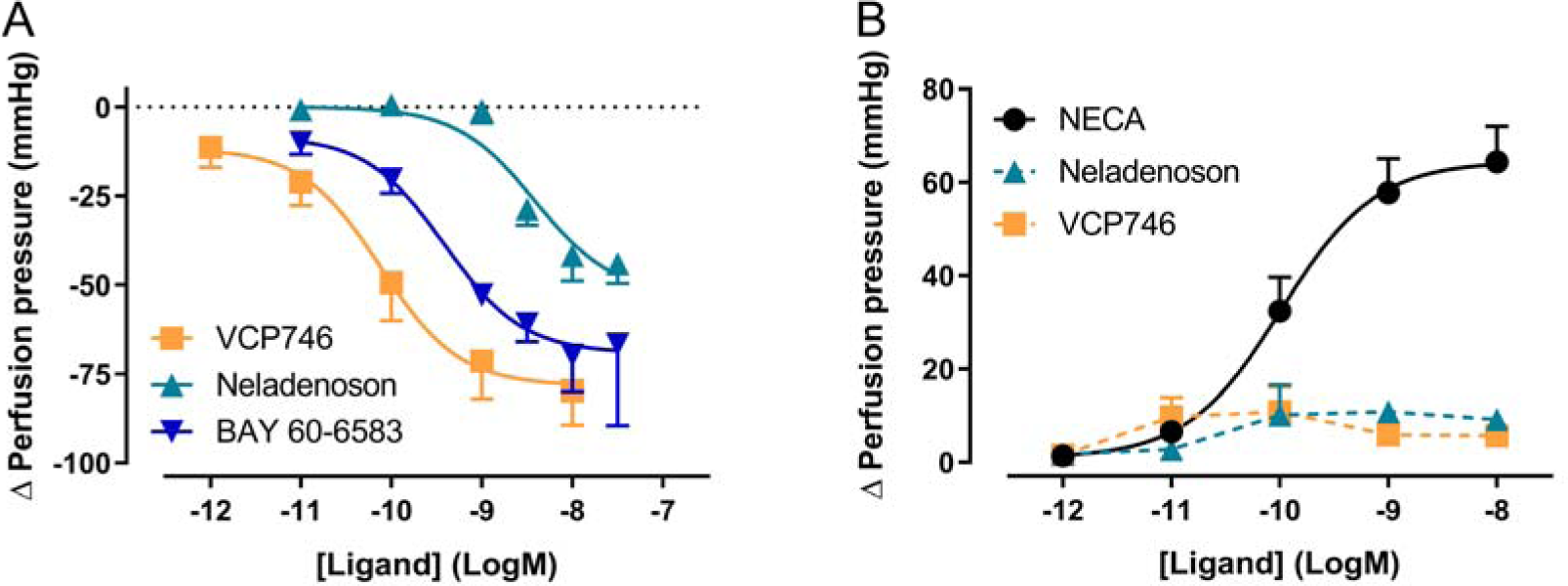
VCP746 and neladenoson induce A_2B_R-mediated renal vasodilation but no A_1_R-mediated vasoconstriction. (A) Renal vasodilation was measured by administering VCP746 or the A_2A_R/A_2B_R agonist BAY 60-6583 to methoxamine-stimulated rat kidney and measuring resulting perfusion pressure (n=3). The concentration-dependent vasodilation produced, with order of potency VCP467 > BAY 60-6583 > neladenson suggest an A_2B_R-mediated mechanism of action, which was confirmed using selective antagonists (Supp Fig 5). (B) In order to measure renal vasoconstriction, kidneys were stimulated with methoxamine/forskolin and then NECA (n=6), neladenoson (n=6) or VCP746 (n=5) was applied. NECA induced a potent constrictor activity, with no response observed to neladenoson/VCP746. This was confirmed to be an A_1_R-mediated effect (Supp Fig 5). Data are expressed as mean ± SEM.

In order to study effects on vasoconstriction, methoxamine-treated kidneys were treated with forskolin to activate adenylate cyclase and reduce vascular tone. Under these conditions, NECA induced renal vasoconstriction (Fig 7B) in a concentration dependent-manner (pEC_50_ = 10.0 ± 0.2, n=6). Similarly, the A_1_R-selective agonist, 2-Me-CCPA, induced a vasoconstrictor response that was abolished by the A_1_R-specific antagonist, SLV320 (Supplementary Fig 5B) indicating that the renal vasodilation is A_1_R-dependent. However, VCP746 and neladenoson had no effect on the vasoconstrictor response (Fig 7B). Since the response is sensitive to A_1_R activation, the lack of effect after VCP746 or neladenoson suggests that the bias profile of these compounds contributes to their lack of renal vasoconstrictor activity.

### Neither VCP746 nor neladenoson induce endothelium-dependent thoracic aorta relaxation

In order to confirm subtype selectivity, VCP746 and neladenoson were tested for effects on rat thoracic aorta relaxation, in the presence and absence of endothelium. NECA relaxed the aorta in an endothelium-dependent A_2A_R-mediated manner (Fig 8; Supplementary Fig 6). In contrast, neither VCP746 nor neladenoson had any effect on the aorta (Fig 8), broadly consistent with their relatively low potency (VCP746) or lack of activity (neladenoson) at the A_2A_R (Fig 2).

**Figure 8.**
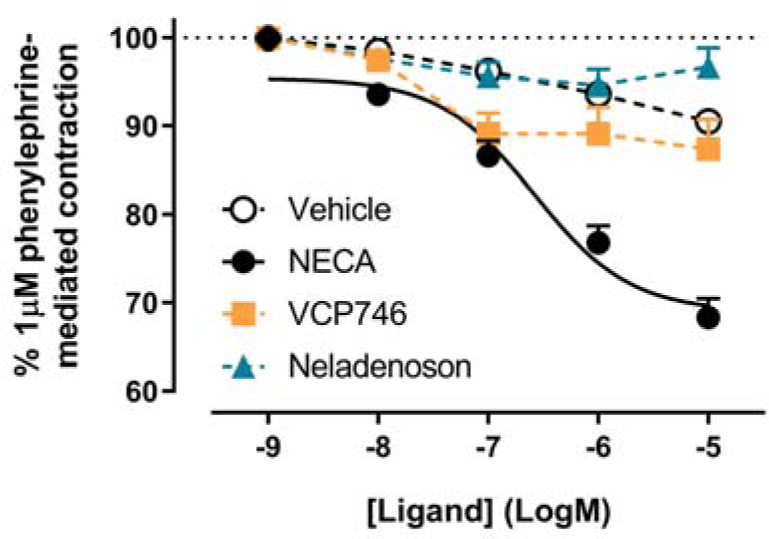
NECA, but not VCP746 or neladenoson induces endothelium-dependent relaxation of rat thoracic aorta. NECA concentration-dependently inhibited phenylephrine-mediated contraction (pEC_50_ 6.6±0.2) while VCP746 and neladenoson displayed negligible effects (n=3). Antagonist studies suggest this is a predominately A_2A_R-mediated mechanism (Supp Fig 6). Data are expressed as mean ± SEM.

## DISCUSSION

The A_1_R has been a focus of heart failure drug discovery efforts for over two decades. The cardioprotective effects of adenosine are well described, but efforts to develop selective A_1_R agonists have been limited by on-target mediated bradycardia, atrioventricular block, and other extra-cardiac undesired effects. Several approaches have been taken to the design of novel agents to mitigate these risks. Bayer developed the A_1_R partial agonists capadenoson and neladenoson (a pro-drug of an active and more soluble metabolite) (*40*), whereas other pre-clinical work identified biased A_1_R agonists that exhibit specific signalling profiles focused on avoiding on-target mediated adverse effects. The A_1_R subtype is a Gα_i/o_-coupled receptor that couples pleiotropically to multiple downstream endpoints, including inhibition of cAMP accumulation, MAPK activation, and increased intracellular calcium mobilisation. Thus, linking a signalling pathway (or subset of pathways) to a specific physiological outcome represents a mechanism whereby agents can be developed that avoid on-target side effects on the kidney, peripheral blood vessels, or the heart.

VCP746 is a tool biased A_1_R agonist that preferentially signals away from intracellular calcium mobilisation compared to inhibition of cAMP accumulation (*32*). Growing evidence points to agonist-induced increases in calcium flux as a predictor of A_1_R-mediated side effects (*9, 32, 37*). We have shown that the A_1_R bias signalling profile of capadenoson, neladenoson and VCP746 are broadly similar in that they are biased away from stimulation of intracellular calcium mobilisation. This phenotype translated to primary cardiomyocytes, isolated rat atria, and *in vivo* in telemetered rats, where both neladenoson and VCP746 had little effect on beat/heart rate, unlike the prototypical adenosine analogue, NECA. Interestingly, at equi-effective concentrations for A_1_R-mediated inhibition of cAMP signalling, neladenoson inhibited the beat rate of ventricular cardiomyocytes and heart rate *in vivo* in contrast to VCP746, suggesting alternative signalling pathways are involved in this process.

Nonetheless, in the clinic, capadenoson and neladenoson had little effect on heart rate in trials in patients with angina pectoris, atrial fibrillation, or congestive heart failure (*42*, *43*; Clinical Trial NLM Identifiers: NCT00568945, NCT03098979, NCT02992288) suggesting that bias in the cAMP-calcium signalling balance downstream of the A_1_R is a critical predictor for avoiding bradycardia (*38, 47*).

A further consideration for A_1_R agonists is their potential for deleterious effects on renal haemodynamics and a number of groups have developed A_1_R *antagonists* for the treatment of impaired renal function in congestive heart failure (*29*). This approach was predicated on stimulating renal vessel relaxation and inhibiting the tubuloglomerular feedback mechanism, thus increasing urine output without worsening glomerular filtration rate. Such a mechanism suggests that A_1_R *agonists* might impair renal function. However, our studies show that VCP746 and neladenoson lack renal vasoconstrictor effects in the methoxamine/forskolin-treated kidney. Again, this indicates that the signalling profile at the A_1_R and/or ancillary pharmacology of these two agents is beneficial when compared to non-selective/non-biased adenosine receptor agonists that cause vasoconstriction. In fact, both neladenoson and VCP746 caused renal vasorelaxation, mediated by the adenosine A_2B_ receptor, which explains the 30-fold discrepancy in potency between the two agents in favour of VCP746. In turn this offers a potential explanation for the small, concentration-dependent increase in heart rate observed with VCP746 *in vivo* that may well be a compensatory response to the increased renal vasorelaxation whilst the rats remain normotensive.

Clinically, neladenoson either had no (*43*) or marginal (*42*) effects on heart rate, a profile largely predicted by pre-clinical models. However, the compound failed to improve cardiac and non-cardiac abnormalities in phase II trials in heart failure patients with either preserved or reduced ejection fraction, and slightly worsened renal function in the latter population (*42, 43*). This clearly differs from the pre-clinical profile, where neladenoson was cardioprotective in a left anterior descending artery occlusion-induced ischaemia model in rats (*40*).

The failure of neladenoson in these clinical trials poses a number of questions for adenosine receptor-targeting heart failure therapeutics. Despite the clinically-proven cardioprotective effects of adenosine (*12, 48, 49*), it might be that selective A_1_R activation, differential A_1_R bias, or targeting patients with chronic, rather than acute, heart failure, could be responsible for the lack of efficacy of neladenoson. Whilst neladenoson has the critical cAMP-calcium bias required for improving therapeutic index, it is also highly biased away from the MAPK pathways, ERK1/2 and Akt1/2/3 phosphorylation. This may be undesirable since ERK1/2 phosphorylation plays a role in cardioprotection (*50–53*), while Akt1/2/3 phosphorylation is generally considered a pro-survival pathway utilised by many cardioprotective agents (*51, 54–56*). It remains to be seen whether therapeutic agents can be developed that have A_1_R cAMP-calcium bias, without the bias away from MAPK signalling. This signalling profile may explain the greater potency of VCP746 in reducing humoral- and inflammation-induced cardiomyocyte hypertrophy compared with neladenoson – herein and (*17*) – despite neladenoson being *more* potent than VCP746 for cAMP inhibition.

With respect to selectivity, our recombinant cell signalling data showed that neladenoson activated neither the A_2A_R or A_3_R subtypes, and (unlike VCP746) is a biased agonist at the A_2B_R (away from MAPK while calcium signalling was undetectable). Activation of A_2B_R reduces fibrosis in heart failure (*21–24*), though the signalling pathway(s) responsible for this are not well defined. Here we showed equivalence for neladenoson and VCP746 for anti-fibrotic efficacy in cardiac fibroblasts, and previous studies have shown that VCP746 exhibits potent A_2B_R-dependent anti-fibrotic activity (*46*). Thus, appropriate A_2B_R signalling profiles for adenosine receptor agonists may be required in heart failure therapeutics. These issues of selectivity and bias are important in the context of treating chronic, as opposed to acute, heart failure, where anti-hypertrophic and anti-fibrotic activity would be desirable. Additionally, the lack of clinical translation calls into question the predictive capacity of rodent models of heart failure (*57*) or their translation to specific stages of the disease in patients.

The concept of biased agonism faces a number of challenges in linking signalling profiles to clinical endpoints and improved therapeutic indices. Despite promising preclinical data, biased agonists have yet to fulfil their full promise in the clinic: the β-arrestin-biased angiotensin AT1 receptor agonist, TRV 027, failed to meet its primary endpoint in a phase IIb study of patients with acute heart failure (*58*). In addition to improving the pre-clinical assessment of what constitutes a biased agonist (e.g. for µ-opioid receptors (*59*)), it is critical to understand the pharmacology of agents that have been developed and clinically-evaluated without reference to the phenomenon of signal bias. In this study we comprehensively compared *in vitro*, *ex vivo*, and *in vivo* properties of the investigational A_1_R agonist, neladenoson, compared to VCP746, a tool A_1_R biased agonist. Retrospective analysis of the signalling profile of neladenoson reinforces the cellular predictor (cAMP-calcium signalling bias) for its lack of effect on heart rate in pre-clinical models and patients. Although the reasons for the lack of clinical efficacy remain unknown, it is possible that fine tuning activity in other pathways (e.g. MAPK) and/or adenosine receptors (e.g. A_2B_R) might represent valid future approaches for the development of novel agents.

## METHODS

### Materials

Research reagents were obtained from the following suppliers: Dulbeco’s modified Eagle medium (DMEM; Life Technologies Australia, 11965118), Hanks’ Balanced Salts (Sigma-Aldrich, H2387), trypsin (Life Technologies Australia, 15090046), antibiotic/antimycotic (Life Technologies Australia, 15240062), penicillin/streptomycin (Gibco, 15140-122), FBS (Gibco), Adenosine deaminase (ADA; Sigma-Aldrich, 10102105001), hygromycin B (Scientific INC, H-1012-PBS), Probenecid (Sigma-Aldrich, P8761), Rolipram (Sigma-Aldrich, R6520), Hoechst33342 (ThermoFisher Scientific, H3570), Propidium iodide (Sigma-Aldrich, P4170), 5-Bromo-2’-deoxyuridine (BrdU; Sigma-Aldrich, B5002), 5’-(N-ethylcarboxamido), 5’-N-ethylcarboxamidoadenosine (NECA; Sigma-Aldrich, E2387), SLV-320 (Tocris, RDS334410), CGS21680 (Tocris, 1063; Sigma-Aldrich, C141), BAY60-6583 (Tocris, 4472), 2’Me-CCPA (Tocris, 2281), MRS1754 (Tocris, 2752), methoxamine (Sigma-Aldrich, M6524), Angiotensin II (AngII; Sigma-Aldrich A9525), interleukin 1beta (IL-1β; R&D systems, 201-LB-005), tumor necrosis factor alpha (TNFα; R&D systems, 210-TA), Lactate dehydrogenase (LDH) Activity Assay Kit (Sigma-Aldrich, MAK066), Adenosine Triphosphate (ATP; Sigma-Aldrich, A26209), Forskolin (Sigma-Aldrich, F3917), Fluo-4 AM (Invitrogen, F14201), Pertussis toxin (PTX; Sigma-Aldrich, P7208), [^3^H]-Leucine (Perkin Elmer, NET135H001MC), [^3^H]-Proline (Perkin Elmer, NET483001MC), Lance cAMP detection kit (PerkinElmer, AD0262) Alphascreen Surefire ERK1/2 (Thr202/Tyr204) Phosphorylation kit (PerkinElmer, TGRESB), Alphascreen Surefire Akt1/2/3 (p-Ser473) Phosphorylation kit (PerkinElmer, TGRA4S), Collagenase Type II (Scimar Australia, LS004176). Neladenoson (as the active metabolite) and capadenoson were synthesised by Servier, and VCP746 was made by SYNthesis Pty. (Melbourne, Australia).

### Cell culture

Flp-IN CHO-A_1_R, -A_2A_R, -A_2B_R, and -A_3_R stable cell lines were generated as previously described (*60, 61*), maintained in DMEM supplemented with 10% FBS and 500 μg/ml hygromycin B and confirmed mycoplasma-free.

### cAMP accumulation

Cells were trypsinised and seeded in DMEM with 10% FBS in 96-well plates at 20,000 cells/well and incubated overnight. Cells were then washed and incubated in cAMP stimulation buffer (140 mM NaCl, 5mM KCl, 0.8 μM MgSO_4_, 0.2 mM Na_2_HPO_4_, 0.44 mM KH_2_PO_4_, 1.3 mM CaCl_2_, 5.6 mM D-glucose, 5 mM HEPES) containing ADA (0.1 U/ml), rolipram (10 μM) and BSA (0.1%) at 37°C in a humidified incubator with 5% CO_2_ for 1h. Compounds were then added and incubated for 30 min. When Gα_i_-mediated signalling was evaluated, 3 μM forskolin was added to the cells 10 min after compound addition. Stimulation was terminated by removal of buffer and replacement with ice-cold 100% ethanol. After ethanol evaporation, cells were lysed in lysis buffer and cAMP levels were detected using the Lance cAMP kit following manufacturer’s instructions. cAMP levels were extrapolated using the standard provided in the kit and then normalised to the forskolin control.

### Calcium mobilisation

Cells were trypsinised and seeded in DMEM with 10% FBS in 96-well plates at 40,000 cells/well for 8h at 37°C in a humidified incubator with 5% CO_2_. Cells were then incubated in serum-free medium overnight, washed and incubated in calcium stimulation buffer (146 mM NaCl, 5 mM KCl, 1mM MgSO_4_, 1.3 mM CaCl_2_, and 1.5 mM NaHCO_3_, 10 mM D-glucose, 10mM HEPES) containing ADA (0.1 U/ml), probenecid (2.5 mM), BSA (0.5%) and Fluo-4 AM (1 μM) for 1 hour. Fluorescence was detected on a FlexStation plate reader (molecular Devices; Sunnyvale, CA, USA) after the automated addition of buffer in the absence or presence of receptor ligands. Data were analysed as the difference between the peak and baseline reads and normalised to the ATP (100 μM) response.

### ERK1/2 and Akt1/2/3 phosphorylation

Cells were trypsinised and seeded in DMEM with 10% FBS in 96-well plates at 40,000 cells/well for 8h at 37°C in a humidified incubator with 5% CO_2_. Cells were then incubated in serum-free medium overnight, and ADA (0.1U/ml) added 1h prior to assay. Cells were then exposed to DMEM in the absence or presence of receptor ligands and agonist concentration-response curves were generated at the time of peak response. Stimulation was terminated by rapid removal of media and addition of 50 μl/well or AlphaScreen SureFire kit lysis buffer. Detection of either ERK1/2 or Akt1/2/3 phosphorylation was performed as described in the corresponding AlphaScreen SureFire kits and fluorescence measured with an EnVision plate reader (PerkinElmer, Boston, MA). Data were normalised to the response elicited upon stimulation of cells with 10% FBS.

### Cell survival

Cells were trypsinised and seeded in DMEM 10% plus FBS in 96-well plates at 40,000 cells/well for 8h at 37°C in a humidified incubator with 5% CO_2_. After 8 hours, plates were rinsed in serum-free DMEM and then incubated in serum-free medium overnight. Media was then changed to fresh, sterile calcium stimulation buffer, containing ADA (0.1U/mL) and pen/strep (1U/mL) and incubated for a further 24h. Hoechst33342 (200 μM; to define all cells) and propidium iodide (PI; 50 μg/ml; to define dead/dying cells) were added to two wells, incubated at 37°C for 30min, and cell nuclei counts detected using the Operetta (PerkinElmer, Boston, MA) using manufacturer’s protocols. This defined 0% cell death for subsequent assay. Varying concentrations of adenosine receptor agonists were added to the remaining wells and cells incubated at 37°C for 24h. Finally, Hoechst33342 and PI stains were added to all wells, incubated for 30 min, and nuclei counts detected on the Operetta. Immediately prior to addition of stains, buffer was removed from two wells and MilliQ water added to lyse cells as a positive control. Data are expressed as a percentage of surviving cells.

### Cardiomyocyte isolation and culture

Neonatal cardiac myocytes (CM) were isolated from 1 to 2 day-old Sprague-Dawley rat pups using enzymatic digestion. Briefly, animals were euthanized and the hearts isolated by thoracic incision and kept in Hanks solution. Then ventricles were isolated and incised at the apex to increase tissue surface exposure. Tissue was then incubated overnight at 4°C with trypsin. After trypsin deactivation with fresh FBS-supplemented DMEM, tissue was further digested by four cycles of collagenase incubation. Cells were then recovered by centrifugation and seeded in DMEM supplemented with FBS on gelatin-coated dishes for 2h in order to select against adherent fibroblasts. The remaining floating cardiomyocytes where then collected, counted, and seeded in either 12-well plates at a density of 300,000 cells/well, or 96-well “chimney well” cell culture plates (Eppendorf) at a density of 37,500 cells/well. Cells were maintained in DMEM supplemented with 10% FBS and BrdU (100 μM) was included for the first three days of culture.

### [^3^H]-Leucine incorporation in cardiomyocytes

To measure hypertrophy by [^3^H]-leucine incorporation, after five days in culture cardiomyocytes were starved overnight in serum-free DMEM and then pre-treated with adenosine receptor agonists or vehicle for 2h before hypertrophic stimuli [IL-1β (10 ng/ml), TNF-α (10 ng/ml), or Ang II (100 nM)], after which 1 μCi of [^3^H]-leucine was added to each well. Cells were incubated at 37°C with 5% CO2 in a humidified incubator for 72h, washed with PBS and lysed using 0.2M NaOH. After adding UltimaGold scintilliant to the samples, radioactivity was detected using a MicroBeta2 Plate Counter (PerkinElmer Life Sciences). Data were normalised to the signal obtained for the vehicle treated samples.

### [^3^H]-Proline incorporation in cardiac fibroblasts

Cardiac fibroblasts were recovered from gelatin-coated dishes described above through trypsination and plated in 12-well plates at a density of 50,000 cells/well for [^3^H]-proline incorporation assays in high glucose DMEM supplemented with 10% FBS. After 4 days cardiac fibroblasts were serum starved in DMEM overnight and then treated with vehicle, VCP746 or neladenoson 2h prior to fibrotic stimuli TGFβ (10 ng/mL) or AngII (10 nM). [^3^H]-Proline (1 μCi/well) was then added to each well. After 72h cells were washed, lysed and radioactivity detected as per [^3^H]-leucine assay, described above.

### Cardiomyocyte LDH release and PI staining

To determine apoptotic effects of adenosine receptor agonists, a combined LDH release and PI stain assay was performed with an identical treatment regimen to [^3^H]-leucine incorporation assays: 2h pre-treatment with adenosine receptor agonists before stimulation with IL-1β (10 ng/ml), TNF-α (10 ng/ml) or Ang II (100 nM) for 72h at 37°C with 5% CO_2_ in a humidified incubator. On the day of assay, 20μL of extracellular media from cardiomyocyte plates was transferred to a new 96-well plate, with NADH standard dilutions, and a colorimetric LDH release assay performed as per manufacturer’s instructions (Sigma-Aldrich; cat no: MAK066). Absorbance (450 nm) was detected at 3 min intervals (FlexStation) until it reached the upper range of standards. Data (“cell viability”) are expressed as a percentage of control-treated wells. In parallel, the cells without washing were treated with DMEM containing Hoescht33342 (200 μM) and PI (50 μg/mL) for 1h at 37°C. Hoechst33342- and PI-positive nuclei were quantified on the Operetta using manufacturer’s protocol. Data (“cell survival”) are a ratio of PI/Hoechst33342 counts, expressed as a percentage of control-treated wells.

### Measurement of beat rate in cardiomyocytes

Rat cardiomyocytes were cultured 4 days in 96-well plates before cells were checked visually to have widespread cell-cell contact and uniform beating across the well. Media was then changed to fresh DMEM with 10% FBS 24h prior to assay, and 0.1U/mL ADA added 2h prior to assay. For assay, spontaneous contractions were brightfield recorded (Nikon Ti-E microscope; 37°C, 5% CO_2_) for 100 frames at 10 frames/sec to define a basal beat rate. Cells were then incubated with ligand for 5 min and recorded again. Quantification of contractions was determined by time-resolved analysis (Image J 1.51n) of peak intensity of several representative cardiomyocytes within the field of view. All data are expressed as % of beat rate prior to addition for each replicate. Where SLV320 (1 μM) was used, it was added 15 min prior to basal recording.

### Measurement of beat rate in isolated rat atria

Experiments were carried out on male Wistar rats (400-450g) from JANVIER Labs (Centre d’Elevage René JANVIER, Le Genest Saint-Isle, France). Rats were anesthetized with sodium pentobarbital (54.7 mg/kg intraperitoneal). Right atria was carefully removed and immediately immersed in physiological salt solution (PSS) at 4°C containing (in mM): NaCl 112, KCl 5, KH_2_PO_4_ 1, MgSO_4_ 1.2, CaCl_2_ 2.5, NaHCO_3_ 29.8, glucose 11.5, EDTA 0.02, pH 7.4±0.05. Preparations were suspended vertically in an organ bath filled with 20 mL of PSS maintained at 35°C and gassed with a mixture of 95% O_2_ + 5% CO_2_. Isometric tension was recorded by means of a force transducer (EMKA Technologies, Paris, France). Atria were stretched to obtain a resting tension of 0.2 to 1 g. Right atria spontaneous beating frequency was measured using specific software (IOX EMKA Technologies, Paris, France). Only preparations with a basal beating rate between 200 and 300 beats per minute (bpm) were included.

After a 30 min equilibration period, dose-response curves were generated with cumulative concentrations (10^−9^M to 10^−5^M) of NECA, CGS21680, BAY60-6583 or 2’MeCCPA every 30 min. Due to their long kinetic of effect, VCP746 and neladenoson were tested with a single concentration (10^−7^M to 10^−5^M) per preparation, beating rate was measured at 120 min after each concentration. To assess NECA specificity, atria were pre-treated (30 min) with the specific A1R antagonist SLV320 at 10nM or 100nM. Then, 2 successive concentrations of NECA (10 and 100 nM) were added for one hour.

Spontaneous beating frequency of right atria, measured at fixed time or concentration, was expressed in bpm. The effects of the compounds were expressed as percent changes from the basal atrial beating frequency.

### Measurement of heart rate in moving, conscious animals

Male Wistar rats (8 weeks old, n=4-7) were implanted with a radio telemetric device equipped with a pressure transducer (HDS10, Data Sciences International) under anesthesia with isoflurane (2%, VETFLURANE®, VIRBAC, France). Rats received buprenorphine (50 µg/kg s.c., BUPRECARE®, Axience SAS, France) for analgesia prior to surgery. After laparotomy the telemetric device catheter was inserted into the abdominal aorta and secured with cellulose patch and tissue adhesive (3M, VETBOND™) around the insertion point. The body of the device was placed in the abdominal cavity and sutured to the inner side of the abdominal musculature then the skin plan was closed.

After a recovery period of two weeks a catheter (polyethylene/silastic) was introduced in jugular vein for drug administration by infusion. Briefly, the jugular vein was dissected and a small incision was made in order to introduce the catheter into the vessel. The catheter was tunnelled subcutaneously to the dorsum of the neck and drawn back up through the skin. Rats were kept on a heating pad (38°C) until fully recovered from anaesthesia. Animals were allowed 2-3 days recovery before being used for the experiment. On the day of study, 20 min infusion protocols were performed at different doses of test compounds to achieve predetermined plasma drug concentrations. Acquisition of heart rate was done with IOX (EMKA, France), sampled and averaged over 5s every 5s. The mean of the values recorded for each dose was calculated using the software Data Analyst (EMKA, France).

### Vasorelaxation of rat thoracic aorta

Experiments were carried out on male Wistar rats (400-450g) from JANVIER Labs (Centre d’Elevage René JANVIER, Le Genest Saint-Isle, France). Rats were anesthetized with sodium pentobarbital (54.7 mg/kg intraperitoneal). The thoracic aorta was quickly removed placed in ice-cold physiological salt solution (PSS) at 4°C containing (in mM): NaCl 112, KCl 5, KH_2_PO_4_ 1, MgSO_4_ 1.2, CaCl_2_ 2.5, NaHCO_3_ 29.8, glucose 11.5, EDTA 0.02, pH 7.4±0.05. Then the aorta was cleaned of adhering fat and connective tissue and cut transversely into 3-4 mm rings denuded or not of endothelium and placed in 20 ml organ baths containing PSS maintained at 37°C with continuous bubbling of 95% O_2_ + 5% CO_2_. Aortic rings were mounted vertically between two stainless wire hooks and then suspended. For isometric force response measurement, the changes in tension of pre-contracted intact or denuded rings were continuously monitored and recorded using specific software (IOX EMKA Technologies, Paris, France). Aortic rings were equilibrated for 60 min with a resting force of 2.5 g.

Aortic rings were constricted with phenylephrine (1 µM) to obtain a steady contraction then relaxed with cumulative acetylcholine concentration (0.01 to 10 µM) to check the integrity of the endothelium. The absence of endothelium was confirmed by the lack of responsiveness to acetylcholine. Aortic rings were washed several times with PSS and equilibrated for 30 min. Then the rings were constricted with phenylephrine (1 µM) to obtain a steady contraction and the agonists were added in cumulated concentrations (1 nM to 10 µM) to the organ bath. At the end of each experiment, the rings were tested for viability by being maximally dilated with 100 µM papaverine. Relaxation was expressed as a percent of maximal relaxation to papaverine and as percent changes from phenylephrine contraction.

### Renal vasodilation and vasoconstriction

Male Wistar rats (300-400 g), purchased from Janvier Labs (Centre d’Elevage René Janvier, Le Genest Saint-Isle, France) were anesthetized intraperitoneal with a mixture of ketamine (110 mg/kg) and xylazine (7.5 mg/kg). The left kidney was exposed by midline ventral laparotomy and the left renal artery was cannulated. The kidney was then perfused at constant flow via a peristaltic pump with warmed (37°C) and oxygenated (95% O_2_-5% CO_2_) Tyrode solution of the following composition (mM): NaCl 137; KCl 2.7; CaCl_2_ 1.8; MgCl_2_ 1.1; NaHCO_3_ 12.0; NaHPO_4_ 0.42; calcium disodium EDTA 0.026 and glucose 5.6. The perfused kidney was removed from the surrounding fat and placed in a perfusion chamber. The change in renal vascular resistance was recorded as changes in renal perfusion pressure (RPP) measured downstream of the pump via a pressure transducer (P10EZ, Statham, France) connected to a data acquisition system (IOX2, EMKA Technologies, France). Pharmacological agents were administered either via an infusion pump placed upstream to the perfusion pump (concentrations expressed below in mol/L) or injected as a bolus of 20 µL (doses expressed in mol) into the perfusion circuit.

In order to study renal vasodilator response, a sustained and stable vasoconstriction was maintained by permanent infusion of methoxamine (10 µmol/L). After stable vasoconstriction, dose-response curves were performed by using BAY60-6583 (1 ρmol to 10 nmol), neladenoson (10 ρmol to 30 nmol), or VCP746 (1 ρmol to 10 nmol). To verify the effect of BAY60-6583 in the renovascular responses to vasodilation, selective A2A (SCH 442416 100nM) or A2B (MRS 1754 100nM) receptor antagonists were administered by perfusion to the kidney 30 min before the agonist. Vasodilation was expressed by the negative delta between the response and Methoxamine constriction.

In order to measure renal vasoconstriction, methoxamine-treated (0.01 µmol/L) kidneys were stimulated with 0.1 µmol/L forskolin. Under these conditions, purine inhibition of adenylate cyclase would blunt the forskolin effect thereby producing vasoconstriction (*62*). Then 2’MeCCPA (1 ρmol to 10 nmol), NECA (1 ρmol to 100 nmol), neladenoson (1 ρmol to 30 nmol) or VCP746 (1 ρmol to 10 nmol) were administered by bolus injection to induce vasoconstriction. Before methoxamine and forskolin infusion two bolus injections of noradrenaline (NA, 0.3 nmol) were performed to obtain reference vasoconstriction. To verify 2’MeCCPA effect on renovascular response, a selective A1 receptors antagonist (SLV320 0.1 and 0.01 µM), was administered 30 min prior agonist administration. Vasoconstriction was expressed as difference between the response and Methoxamine/Forskolin induced constriction, or as the % of NA constriction for the experiment with 2’MECCPA.

### Statistics

All data are expressed as mean ± SEM or mean ± SD (as indicated) of *n* experiments. For concentration response data, curves were analysed using three-parameter nonlinear curve fitting (GraphPad Prism 8.02) of grouped data. pK_b_ values of VCP746 at hA_3_ receptor were determined using the Schild method (*63*) and calculated using the operational model plus agonism (GraphPad Prism 8.02). Statistical analysis was performed using paired student’s t-test, repeated measures one-way analysis of variance (ANOVA) or two-way ANOVA with Dunnett’s post-hoc test, as indicated in results. p<0.05 was considered significant.

Ligand bias at adenosine receptors was analysed using methods described previously (*31, 64*). Cyclic AMP accumulation, calcium mobilisation and protein phosphorylation assays were performed using all four AR agonists in parallel, allowing calculation of Log(τ/K_A_) values for each individual *n*, and subsequent calculation of bias to include *weighted* mean±SEM of *n*. Bias (Log(τ/K_A_), ΔLog(τ/K_A_), ΔΔLog(τ/K_A_)) was calculated in Microsoft Excel (2010) and bias plots generated using the radar plot feature. Analysis of ΔLog(τ/K_A_) and ΔΔLog(τ/K_A_) were performed using multiple two-way ANOVA with multiple comparisons (GraphPad Prism 8.02).

### Study approval

Animal experiments were conducted in accordance to either Servier Ethical committee guidelines or the Monash Institute of Pharmaceutical Sciences animal ethics committee-approved protocols (ethics approval number: MIPS.2017.18) and conformed to the requirements of the National Health and Medical Research Council of Australia *Code of practice for the care and use of animals for scientific purposes*.

## ACKNOWLEDGMENTS

The authors would like to thank Dr Joanne Baltos, Dr Elizabeth Vecchio and Dr Anh Nguyen for gift of the CHO cells.

## AUTHORS CONTRIBUTIONS

PR and JM performed all signalling assays, and all experiments in isolated cardiomyocytes and cardiac fibroblasts and all subsequent analysis. SC performed in vivo experiments. JP performed isolated aorta, kidney and atria experiments. PR, JM and CJL prepared the manuscript and PMS, SC and JP provided feedback on the manuscript. PJW, ACS, WNC, RJS, PMS, LTM, SC, JP and MF provided feedback throughout the project.

## SUPPLEMENTARY TABLES

**Supp table 1.** Potencies (pEC_50_) and E_max_/Span of responses at A_1_R (relative to indicated control) of the adenosinergic compounds from data in Figure 1. Data are mean ± SEM and experimental *n* are indicated in brackets.

| Pathway/<br>Ligand | cAMP inhibition<br>(6) |  | Ca<sup>2+</sup><sub>i</sub><br>(4) |  | pERK1/2<br>(5) |  | pAKT1/2/3<br>(3) |  | Cell Survival<br>(3) |  |
| --- | --- | --- | --- | --- | --- | --- | --- | --- | --- | --- |
|  | pEC<sub>50</sub> | E<sub>max</sub><br>(%Fsk) | pEC<sub>50</sub> | E<sub>max</sub><br>(%ATP) | pEC<sub>50</sub> | E<sub>max</sub><br>(%FBS) | pEC<sub>50</sub> | E<sub>max</sub><br>(%FBS) | pEC<sub>50</sub> | Span<br>(%Basal) |
| NECA | 8.8 ± 0.1 | 13.5 ± 2.2 | 7.6 ± 0.1 | 38.0 ± 1.2 | 9.1 ± 0.1 | 87 ± 4.1 | 8.5 ± 0.1 | 109 ± 4 | 9.2 ± 0.2 | 21.6 ± 1.8 |
| VCP746 | 7.7 ± 0.1 | 13.4 ± 3.2 | 6.8 ± 0.5 | 8.1 ± 1.3 | 8.2 ± 0.1 | 105 ± 3 | 7.7 ± 0.1 | 91 ± 5 | 8.3 ± 0.3 | 21.1 ± 3.1 |
| Capadenoson | 9.5 ± 0.1 | 16.5 ± 2.1 | 7.6 ± 0.1 | 19.2 ± 0.9 | 8.9 ± 0.1 | 102 ± 4 | 8.5 ± 0.1 | 93 ± 3 | 9.5 ± 0.2 | 22.5 ± 1.7 |
| Neladenoson | 8.9 ± 0.1 | 13.2 ± 2.7 | 7.2 ± 0.2 | 16.6 ± 1.1 | 7.1 ± 0.1 | 117 ± 4 | 6.6 ± 0.1 | 80 ± 5 | 9.0 ± 0.2 | 23.9 ± 1.9 |

**Supp table 2.** Potencies (pEC_50_) and E_max_ responses at A_2A_R (relative to indicated control) of the adenosinergic compounds from data in Figure 2. “∼” indicates true E_max_ (and therefore potency) could only be estimated, and “----” indicates no response was observed. Data are mean ± SEM and experimental *n* are indicated in brackets.

| Pathway/<br>Ligand | cAMP stimulation<br>(3) |  | Ca<sup>2+</sup><sub>i</sub><br>(3) |  | pERK1/2<br>(3) |  | pAKT1/2/3<br>(4) |  |
| --- | --- | --- | --- | --- | --- | --- | --- | --- |
|  | pEC<sub>50</sub> | E<sub>max</sub><br>(%Fsk) | pEC<sub>50</sub> | E<sub>max</sub><br>(%ATP) | pEC<sub>50</sub> | E<sub>max</sub><br>(%FBS) | pEC<sub>50</sub> | E<sub>max</sub><br>(%FBS) |
| NECA | 7.8 ± 0.1 | 17.5 ± .5 | ---- | ---- | 8.7 ± 0.2 | 35.4 ± 2 | 8.4 ± 0.3 | 6.7 ± .7 |
| VCP746 | ~5.5 | ~15.6 | ---- | ---- | 7.5 ± 0.2 | 25.0 ± 2 | 7.0 ± 0.3 | 4.8 ± 0.6 |
| Capadenoson | ~5.4 | ~15.3 | ---- | ---- | 6.7 ± 0.1 | 31.3 ± 2 | 6.5 ± 0.3 | 5.8 ± 0.8 |
| Neladenoson | ---- | ---- | ---- | ---- | ---- | ---- | ---- | ---- |

**Supp table 3.**
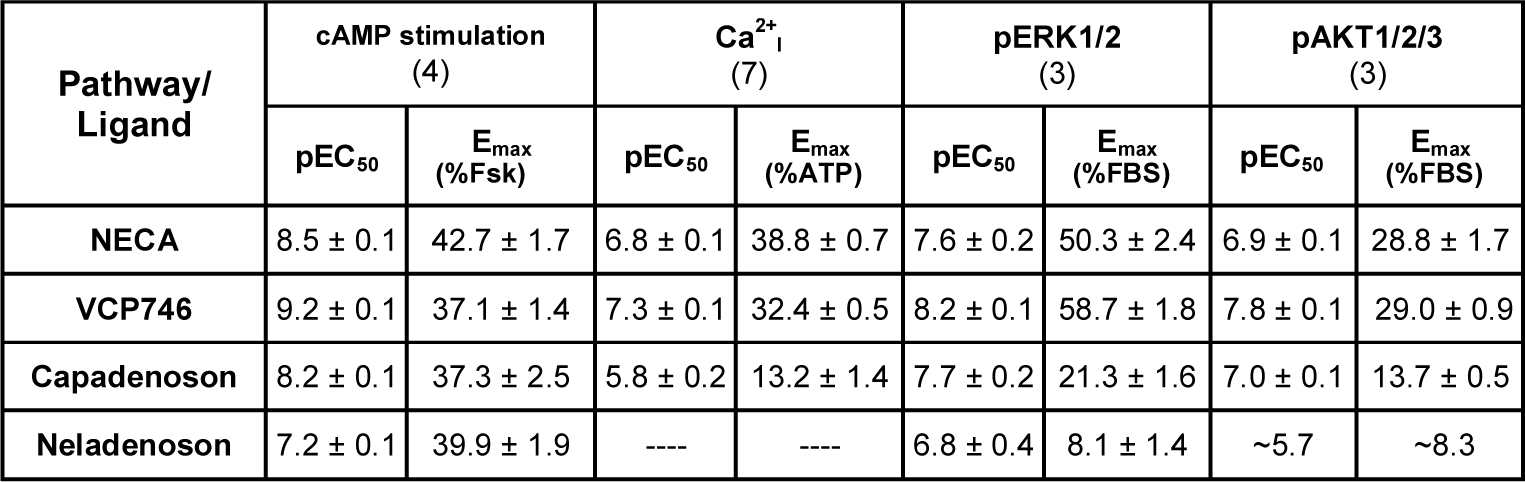
Potencies (pEC_50_) and E_max_ responses at A_2B_R (relative to indicated control) of the adenosinergic compounds from data in Figure 2. “∼” indicates true E_max_ (and therefore potency) could only be estimated, and “----” indicates no response was observed. Data are mean ± SEM and experimental *n* are indicated in brackets.

| Pathway/<br>Ligand | cAMP stimulation<br>(4) |  | Ca<sup>2+</sup><sub>i</sub><br>(7) |  | pERK1/2<br>(3) |  | pAKT1/2/3<br>(3) |  |
| --- | --- | --- | --- | --- | --- | --- | --- | --- |
|  | pEC<sub>50</sub> | E<sub>max</sub><br>(%Fsk) | pEC<sub>50</sub> | E<sub>max</sub><br>(%ATP) | pEC<sub>50</sub> | E<sub>max</sub><br>(%FBS) | pEC<sub>50</sub> | E<sub>max</sub><br>(%FBS) |
| NECA | 8.5 ± 0.1 | 42.7 ± 1.7 | 6.8 ± 0.1 | 38.8 ± 0.7 | 7.6 ± 0.2 | 50.3 ± 2.4 | 6.9 ± 0.1 | 28.8 ± 1.7 |
| VCP746 | 9.2 ± 0.1 | 37.1 ± 1.4 | 7.3 ± 0.1 | 32.4 ± 0.5 | 8.2 ± 0.1 | 58.7 ± 1.8 | 7.8 ± 0.1 | 29.0 ± 0.9 |
| Capadenoson | 8.2 ± 0.1 | 37.3 ± 2.5 | 5.8 ± 0.2 | 13.2 ± 1.4 | 7.7 ± 0.2 | 21.3 ± 1.6 | 7.0 ± 0.1 | 13.7 ± 0.5 |
| Neladenoson | 7.2 ± 0.1 | 39.9 ± 1.9 | ---- | ---- | 6.8 ± 0.4 | 8.1 ± 1.4 | ~5.7 | ~8.3 |

**Supp table 4.**
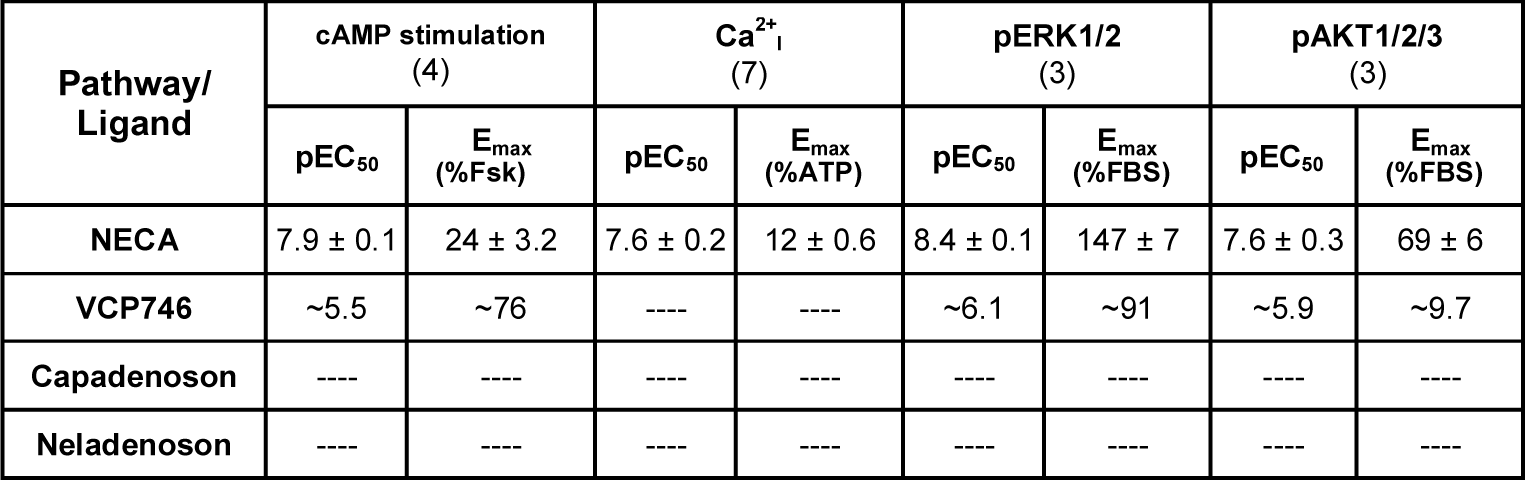
Potencies (pEC_50_) and E_max_ responses at A_3_R (relative to indicated control) of the adenosinergic compounds from data in Figure 2. “∼” indicates true E_max_ (and therefore potency) could only be estimated, and “----” indicates no response was observed. Data are mean ± SEM and experimental *n* are indicated in brackets.

| Pathway/<br>Ligand | cAMP stimulation<br>(4) |  | Ca<sup>2+</sup><sub>i</sub><br>(7) |  | pERK1/2<br>(3) |  | pAKT1/2/3<br>(3) |  |
| --- | --- | --- | --- | --- | --- | --- | --- | --- |
|  | pEC<sub>50</sub> | E<sub>max</sub><br>(%Fsk) | pEC<sub>50</sub> | E<sub>max</sub><br>(%ATP) | pEC<sub>50</sub> | E<sub>max</sub><br>(%FBS) | pEC<sub>50</sub> | E<sub>max</sub><br>(%FBS) |
| NECA | 7.9 ± 0.1 | 24 ± 3.2 | 7.6 ± 0.2 | 12 ± 0.6 | 8.4 ± 0.1 | 147 ± 7 | 7.6 ± 0.3 | 69 ± 6 |
| VCP746 | ~5.5 | ~76 | ---- | ---- | ~6.1 | ~91 | ~5.9 | ~9.7 |
| Capadenoson | ---- | ---- | ---- | ---- | ---- | ---- | ---- | ---- |
| Neladenoson | ---- | ---- | ---- | ---- | ---- | ---- | ---- | ---- |

## SUPPLEMENTARY FIGURES

**Supp Figure 1.**
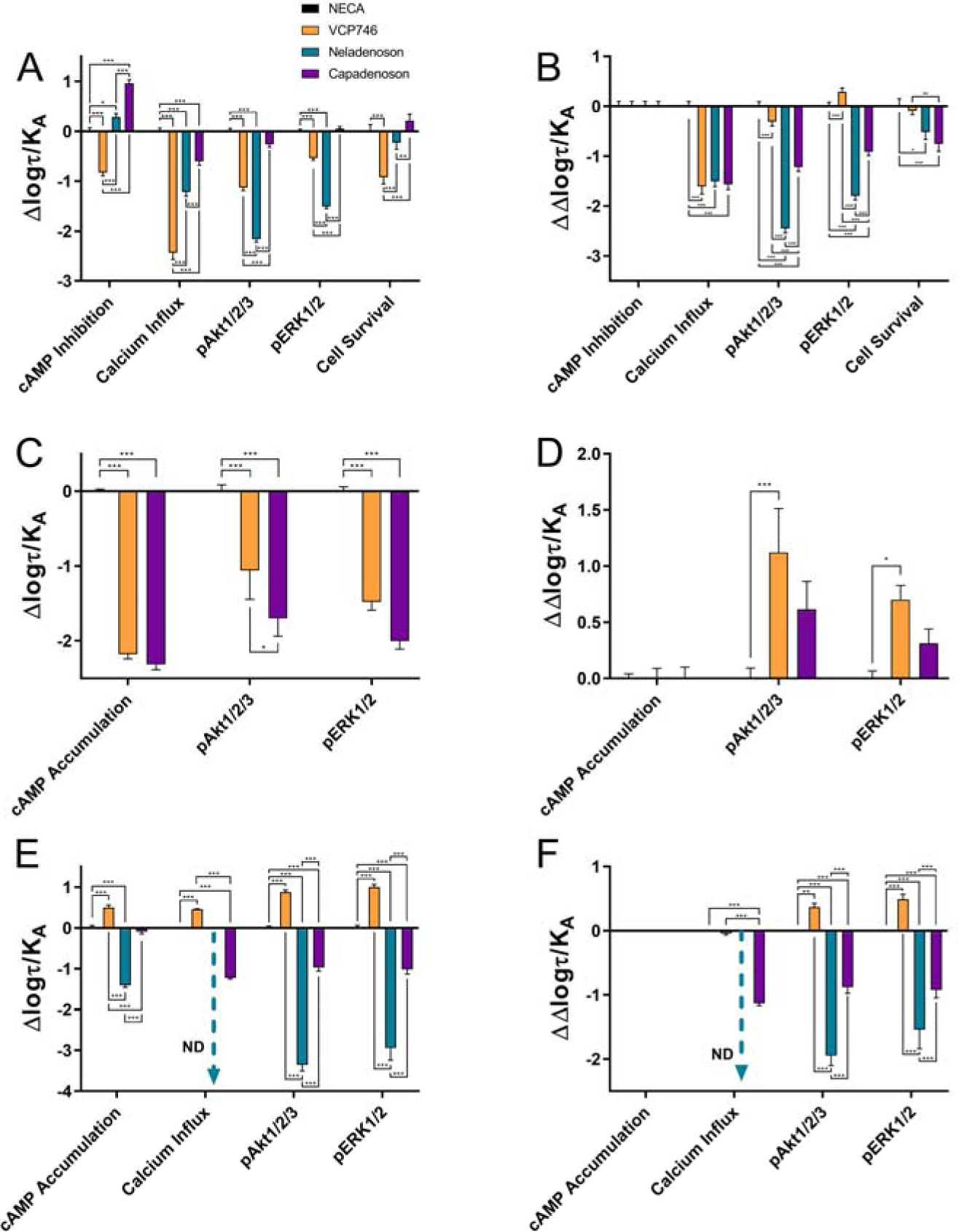
Bias data were normalised to NECA within each cell line (A, C and E), before normalisation to the canonical Gi/Gs pathway cAMP (B, D and F). Each experiment was performed with all ligands so that bias could be calculated within each *n*. Calcium influx bias of neladenoson at A_2B_ (E and F) was not determined (*ND*) due to the lack of calcium response. As calculated bias from each experiment contained error, data are expressed as *weighted* mean ± SEM, and therefore contain no individual data points. The weighted mean 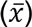 is calculated by:

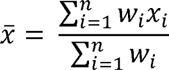

where *w_i_* is the weight of each experiment and *x_i_* is the mean from each experiment. The weighting of each experiment (*w_i_*) is calculated by:

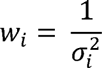

where 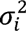 is the variance (square of the standard deviation) from each experiment. The error (SEM) is propagated through by dividing 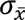 (standard deviation of the weighted mean) by *n*, where:

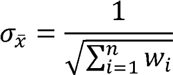

Data for each pathway were analysed by two-way ANOVA, *(p<0.05), **(p<0.01), ***(p<0.001).

**Supp Figure 2.**
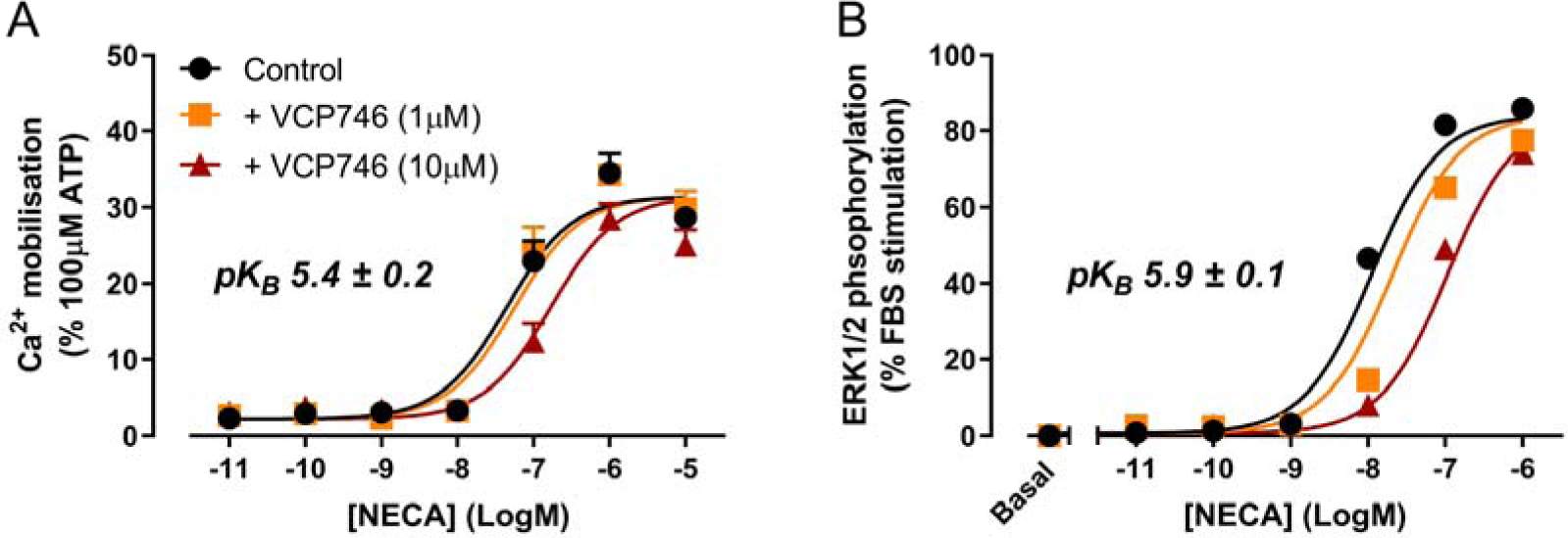
VCP746 acts as a low affinity antagonist on the A_3_R. VCP746 (1μM or 10μM) was applied 15 minutes prior to NECA in calcium influx (A; n=4) and ERK1/2 phosphorylation assays (B; n=4). pK_B_ values were calculated using the Gaddum/Schild EC_50_ shift method in GraphPad Prism 8.02.

**Supp Figure 3.**
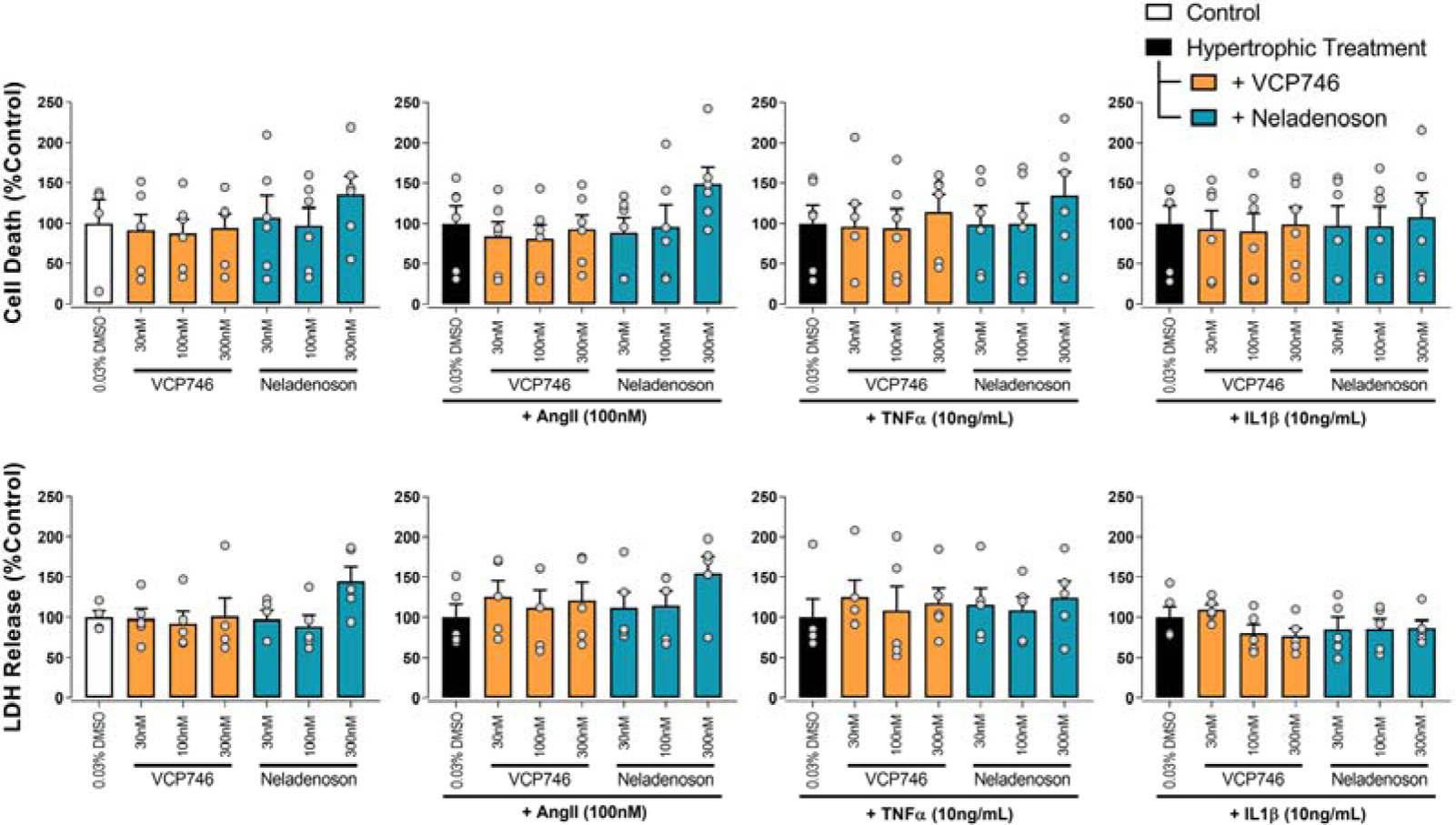
Adenosinergic treatments did not affect NVCM cell viability. Rat primary neonatal ventricular cardiomyocytes treated identically to hypertrophy assays (Fig 4) were analysed for effects on viability through two methods: proportion of cells stained by PI relative to the entire population determined by Hoescht33324 (top panels; n=5); and the release of LDH from cells into the media as an indication of membrane permeability (lower panels; n=5). In both cases data are normalised to control within that hypertrophic treatment. No statistical significance was determined (repeated measures one-way ANOVA with Dunnett’s post-test, compared to control or hypertrophic treatment alone).

**Supp Figure 4.**
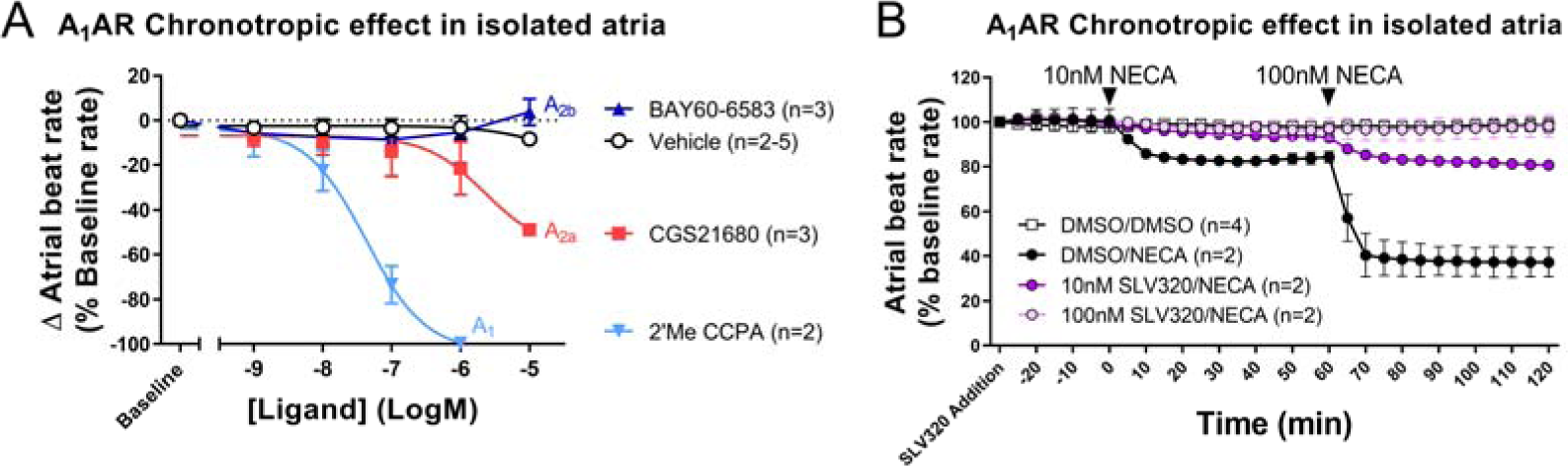
Ex vivo chronotropic effects are A_1_R-mediated. (A) In isolated atria, the effect of NECA was recapitulated with the A_1_R agonist 2’Me CCPA, and to a lesser extent the A_2A_R-selective agonist CGS21680 (although this may be via activity at A_1_R). (B) Pretreatment with the A_1_R antagonist SLV320 inhibits NECA negative chronotropic response in isolated atria in a concentration-dependent manner. Data are displayed as mean ± SD.

**Supp Figure 5.**
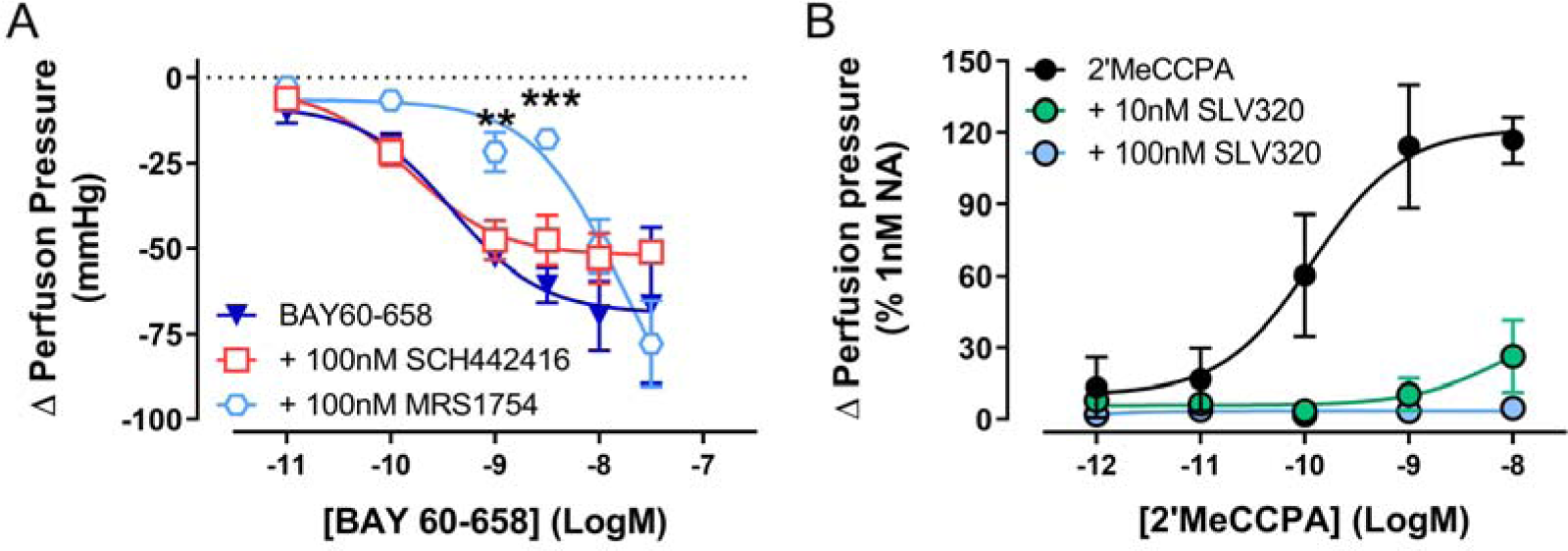
Ex vivo renal vasodilation or vasoconstriction are A_2B_R- or A_1_R-mediated, respectively. (A) The A_2B_R-agonist BAY 60-658 (n=3) reduced perfusion pressure, which was inhibited by pretreatment with the A_2B_R-selective antagonist MRS1754 (n=3; two-way ANOVA, Dunnett’s post-test), but not the A_2A_R-selective antagonist SCH442416 (n=3). Data are displayed as mean ± SEM. 2’MeCCPA (n=3) promoted an increase in perfusion pressure (B), which was entirely A_1_R-mediated as it was concentration-dependently inhibited by pretreatment with the A_1_R-selective antagonist SLV320 at 10nM (n=4), and 100nM (n=2). Data are displayed as mean ± SD.

**Supp Figure 6.**
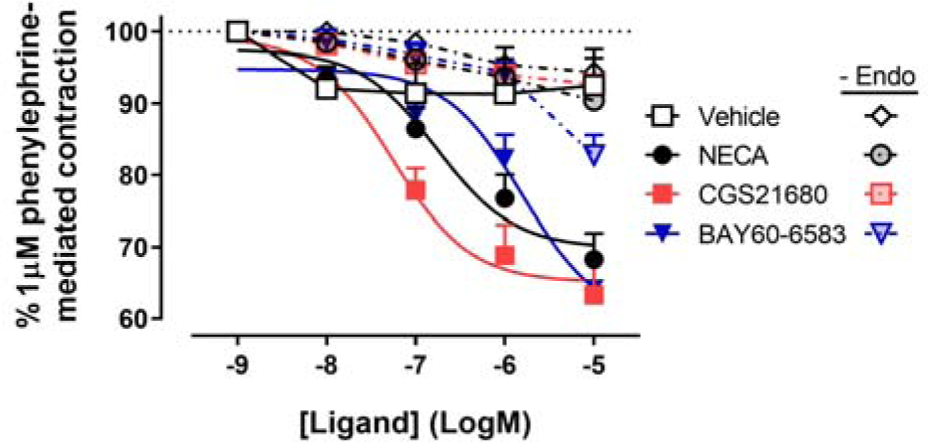
*Ex vivo* rat thoracic aorta relaxation effects are dependent on endothelium and occur through activation of A_2A_R and A_2B_R. (A) Like NECA (pEC_50_ 6.8±0.2, n=9) the A_2A_R-selective agonist CGS21680 (pEC_50_ 7.3±0.2, n=8) and the A_2B_R-selective agonist BAY 60-6583 (pEC_50_ 5.8±0.2, n=10) produced concentration-dependent inhibition of phenylephrine-mediated aorta relaxation. Removing the endothelium completely abrogated the response to both CGS21680 (n=8) and BAY60-6583 (n=5), indicating this A_2_R-mediated relaxation effect is an endothelium-dependent mechanism. Data are expressed as mean ± SEM.

